# Episodic memory ERPs reflect preclinical Alzheimer’s disease progression

**DOI:** 10.64898/2026.07.11.737503

**Authors:** Filipa Raposo Pereira, Maximilien Chaumon, Bruno Dubois, Hovagim Bakardjian, Katia Andrade, Marie-Odile Habert, Nadjia Younsi, Valentina La Corte, Nathalie George, INSIGHT-preAD study group

## Abstract

**Background:** Alzheimer’s disease (AD) neuropathology emerges decades before symptoms, requiring sensitive, non-invasive markers for early diagnosis. We investigated behavioural and EEG markers of episodic memory in cognitively normal older adults at-risk for AD.

**Methods:** Forty-five INSIGHT-preAD cohort participants (15 progressors to prodromal AD, 15 stable β-amyloid positive [Aβ+] participants, 15 controls [Aβ-]) completed an Old/New word recognition task during high-density EEG. Behavioural performance and memory-related evoked-related potentials (ERPs; P3, FN400, P600, post-retrieval monitoring) were analysed longitudinally over 5 years.

**Results:** Progressors showed significantly reduced accuracy, sensitivity, and slower responses. They showed attenuated FN400, P600, and monitoring amplitudes over left parieto-occipital regions. Source analysis revealed hypoactivation in the left inferior frontal and right posterior cingulate cortex, across components with additional disruption during P600. Stable Aβ+ individuals showed milder P600 hypoactivations.

**Discussion:** Behavioural and ERP alterations preceded clinical decline, supporting EEG-based markers of episodic memory decline for preclinical AD risk stratification.

## 1. Background

A new case of Alzheimer’s disease (AD) or related dementia arises every 3.2 seconds worldwide, yet approximately 75% remain undiagnosed (Long et al., 2023; Alzheimer’s Association, 2024). This highlights the urgency for cost-effective, simple, and minimally invasive diagnostic tools to tackle the growing global aging challenge (Dubois et al., 2018; Long et al., 2023).

Episodic memory (EM) alterations—difficulties in retrieving personal events with their spatiotemporal context—are a defining feature of typical AD, often preceded by sustained subjective memory complaints (SMC) (Grober et al., 2000; Grober et al., 2018; Dubois et al., 2018; Jessen et al., 2020; Raposo Pereira et al., 2024a). However, neuropathological hallmarks might begin 20 years before symptoms, with amyloid-β (Aβ) plaques and tau tangles gradually accumulating in the medial temporal, frontal, and parietal lobes (TL, FL, PL, respectively) (Braak and Braak, 1991; Jack et al., 2010; Busche and Hyman, 2020; Hampel et al., 2021; Zamani et al., 2025).

Their accumulation is involved in neuronal death and synaptic dysfunction, potentially desynchronising large-scale memory networks, subtending AD amnesic phenotype (Olichney et al., 2011; Musaeus et al., 2019; Babiloni et al., 2020).

Word list-learning neuropsychological tests, such as the Free and Cued Selective Reminding Test (FCSRT), can be used to probe EM by requiring recognition of previously encoded (“Old”) among novel (“New”) words, engaging familiarity and recollection-based retrieval (Grober et al., 1988; Grober et al., 2000; Rugg and Curran, 2007; Dubois et al., 2018; Raposo Pereira et al., 2024a). Their sensitivity to subtle decline depends on consolidation delays, as longer intervals weaken familiarity and increase reliance on recollection (Rugg and Curran, 2007). When combined with electroencephalography (EEG), Old/New word recognition tasks enable investigating memory-related neural dynamics through event-related potentials (ERPs) in response to Old and New words—i.e., the old/new effect (Picton et al., 2000; Friedman, 2013). Variations in ERP latency, amplitude, and scalp topography offer insight into distinct EM processes, constituting potential early AD biomarkers (Friedman, 2013; Paitel et al., 2021).

Four ERP components consistently associated to EM—P3 (250–350 ms), FN400 (400–580 ms), P600 (600–800 ms), and post-retrieval monitoring ERP (800–1200 ms)—are candidate markers of AD-related memory impairment (Tsivilis et al., 2015; Musaeus et al., 2019; Paitel et al., 2021). The P3, typically distributed over left prefrontal and mid-parietal regions during correctly recognised words, reflects goal-directed attention and encoding, but lacks diagnostic specificity (Horváth et al., 2018; Musaeus et al., 2019). In contrast, the classic dual-retrieval FN400/P600 components provide greater AD-specificity. The FN400, a fronto-central negative deflection sensitive to novelty and repetition, indexes context-free familiarity (“knowing”), whereas the later P600, a centro-parietal positivity sensitive to encoding depth and retrieval, indexes context-dependent recollection (“remembering”) (Olichney et al., 2008; Olichney et al., 2011; Friedman, 2013; Tsivilis et al., 2015). During AD progression, both often show reduced amplitude and delayed latency, FN400 mainly during mild AD—reflecting impaired access to information, while P600 alteration appears also preclinically, consistent with early sensory and contextual retrieval dysfunction (Olichney et al., 2013; Paitel et al., 2021). A later prolonged, right fronto-central, post-retrieval monitoring component, associated with response evaluation and error monitoring, tends to show increased amplitude and asymmetry in AD, interpreted as reduced confidence and greater reliance on guessing (Roper, 2011; Razafimahatratra et al., 2023).

Recent reviews and clinical studies further validate ERP–behavioural approaches as sensitive, non-invasive, cost-effective tools capable of detecting subtle cognitive decline and predict progression among at-risk individuals, reducing underdiagnosis (Olichney et al., 2011; Friedman, 2013; Cecchi et al., 2015; Musaeus et al., 2019; Paitel et al., 2021; Liang et al., 2025). However, EEG remains underused in AD risk assessment.

We used an Old/New word recognition paradigm based on the FCSRT verbal content with an extended consolidation interval, to investigate early ERP and behavioural markers of EM decline predictive of AD within the monocentric and multidisciplinary INSIGHT-preAD longitudinal cohort (Dubois et al., 2018). We retrospectively evaluated the task sensitivity to ERP (P3, FN400, P600, post-retrieval monitoring) and behavioural (accuracy, reaction time [RT], d-prime [d′], criterion) changes associated with subtle AD-related EM alterations over 5 years (M0–M60) in cognitively normal older adults with persistent SMCs. We assessed the impact of Aβ burden and clinical progression to AD considering 15 participants who progressed to prodromal AD, 15 matched participants with Aβ burden (Aβ+) who remained cognitively stable during follow-up, and 15 control participants with below-threshold Aβ burden (Aβ-).

## 2. Methods

### 2.1 Participants

The INSIGHT-preAD cohort comprises 318 cognitively normal older adults at-risk to AD, with subjective memory complaints (SMC) for at least 6 months (McNair test) and with/without the presence of Aβ burden (Aβ) and/or neurodegeneration (N) biomarkers (McNair and Kahn, 1983; Dubois et al., 2018). They were followed-up yearly for five years (M0 to M60) to monitor their risk of progression towards AD. Inclusion criteria included: 70–85 years of age; mini-mental state examination (MMSE) total score ≥27/30; FCSRT total recall (TR) score ≥44/48; clinical dementia rating (CDR) of 0; adequate sensory acuity (Folstein et al., 1975; Morris, 1993; Grober et al., 2000; Dubois et al., 2018; Grober et al., 2018). Exclusion criteria included: under guardianship or institutionalisation; illiteracy; AD or any other neurological diagnoses; MRI incompatibility (Dubois et al., 2018). Cognitive decline (in MMSE, CDR and/or FCSRT-TR) in two consecutive 6-month assessments prompted independent re-assessment by two neurologists, a neuropsychologist, and a neuroimaging expert (Dubois et al., 2018; Habert et al., 2018). In case of medical diagnosis confirmation or death, participation in the study was ceased. After presenting study conditions, all participants provided informed consent and willingness to commit to the study duration; study drop-out was possible at any time.

During follow-up, 15 individuals progressed to prodromal AD (progressors), an early symptomatic phase defined by AD’s clinical phenotype—the amnestic hippocampal syndrome (AHS)—and at least one AD biomarker (Dubois et al., 2018; Raposo Pereira et al., 2024b). Progressors were Aβ+ (8 Aβ+/N- and 7 Aβ+/N+; see below for Aβ and neurodegeneration [N] cutoffs), and were included only until diagnosis as follows: M18 (n=1 diagnosed), M24 (n=3), M36 (n=1), M42 (n=1), and M60 (n=9). As an exploratory approach, we selected two groups (n=15 each) of cognitively normal elderly individuals with SMC who remained free of diagnosis during follow-up: A control group (Aβ-/N-) and a stable Aβ-positive group (8 Aβ+/N- and 7 Aβ+/N+).^7^ These groups were matched with the progressors on age, sex, education, and—as much as possible—on their Aβ burden and neurodegeneration levels. This design enabled us to assess the effect of Aβ-burden (controls vs. stable) and disease progression (stable vs. progressors).

### 2.2 Ethics

This study followed the guidelines of the French law n° 2004-806 (9/08/2004), the Good Clinical Practice principles (I.C.H v.4 of 1/05/1996, decision of 24/11/2006), the Helsinki Declaration (Ethical Principles for Medical Research involving Human Subjects, Tokyo, 2004), and the human ERP research criteria. The Pitié-Salpêtrière Hospital ethics committee endorsed the IN-SIGHT-PreAD study protocol (IDRCB: 2012-A01731-42), and the INSIGHT-preAD scientific committee endorsed the current study. All the data from the INSIGHT-cohort was collected by certified health staff qualified to do so.

### 2.3 Procedure

The INSIGHT-preAD highly multimodal data were collected monocentrically at the Institute for Memory and Alzheimer’s Disease (IM2A), Pitié-Salpêtrière Hospital, Paris. In this study, we used a subset of the data. This included: FDG-PET (neurodegeneration proxy), Aβ-PET, and structural MRI collected at baseline; neuropsychological data (MMSE, FAB, McNair, FCSRT), EEG and associated behavioural data from the Old/New word recognition task, collected yearly from M0 to M60 (Dubois et al., 2018). These assessments were performed on the same day, with imaging and neuropsychological testing in the morning, and EEG recording in the afternoon, resulting in a 4-6 hour interval between the FCSRT word list encoding and the EEG Old/New recognition task that used FCSRT words as Old words (Dubois et al., 2018; Raposo Pereira et al., 2024b). After electrode placement (10-20 min), each EEG session comprised: resting-state (2 min); familiarisation task (presentation of a 6-words sentence followed by their recognition among distractors); Old/New word recognition task (∼20 min); resting-state (2min).

### 2.4 Neuropsychological Assessments

The French socio-cultural scale (*Niveau socio-culturel*, NSC) ranging from 1 (primary education) to 8 (higher education, e.g., PhD), indexed education level. Cognitive functioning was assessed using the 30-point MMSE, covering: orientation (time and place); memory; attention and problem solving; language; and visual-spatial skills (Folstein et al., 1975). Total score <27 prompts further evaluation (Folstein et al., 1975; Dubois et al., 2018). Executive functioning was evaluated via the 18-point FAB (0–3 each), covering: conceptualisation and abstract reasoning; mental flexibility; interference sensitivity; inhibitory control; motor programming; and environmental autonomy. Total score ≤12 suggests AD profile (Dubois et al., 2000). SMC were measured using the 15-item McNair Frequency of Forgetting Questionnaire (rarely to always), completed by participants and companions. Scores ≥15 indicate an abnormal SMC level (McNair and Kahn, 1983).

EM was quantified by the 16-points FCSRT French version (Grober et al., 1988; Grober et al., 2000; Grober et al., 2018; Van der Linden et al., 2004). The test includes a controlled encoding and learning phase where participants view four cards (four words each) and identify targets cued by orally presented semantic categories (e.g., judo-sport). This was followed by immediate recall (IR) for the same items, and cued recall for missed items until they were retrieved. Immediate and cued recall blocks were repeated three times and interleaved with a 20-second interference task (backwards counting). After 20 minutes, delayed free recall (DFR) and delayed cued recall were performed. EM performance was assessed by free recall (FR; sum of IR trials; 0–48) and total recall (TR; sum of IR & cued recall trials; 0–48), with TR ≤41 suggesting possible AD-related impairment (Grober et al., 2000; Dubois et al., 2018).

### 2.5 PET acquisitions

A Philips Gemini GXL CR-PET scanner was used to acquire a standardised uptake value ratio (SUVr) of Aβ at session M0, 50 min after an injection of 370 MBq (10 mCi) ¹⁸F-florbetapir. The Aβ+ vs. Aβ– status was defined at a SUVr cut-off >0.7918 based on the mean uptake in cortical regions of interest (i.e., bilateral precuneus, posterior/anterior cingulum, parietal & temporal associative cortices and orbitofrontal cortex) relative to a reference region (i.e., whole cerebellum plus pons) following CAEN linear conversion method calculated for the INSIGHT-preAD study (Dubois et al., 2018; Habert et al., 2018).

Hypometabolism (low glucose uptake) was considered as an expression of Neurodegeneration (N) quantified in fluorodeoxyglucose (FDG) SUVr collected after an injection of 2 MBq/kg ^18^FFDG (30min). The N + vs. N– status was defined at a FDG-SUVr cut-off <2.27 obtained through the SUVr mean uptake of cortical regions of interest (i.e., posterior cingulate cortex, inferior parietal lobe, precuneus, inferior temporal gyrus) relative to a reference region (pons), similar as above (Jack et al., 2017; Dubois et al., 2018).

### 2.6 Structural MRI acquisition (sMRI) and segmentation

The sMRI data was acquired with a 3T high-resolution Tesla scanner (Siemens Magnetom Verio, Siemens Medical Solutions, Erlangen, Germany) comprising a quadrature-detection head coil with 12 channels (transmit-receive circularly polarised CP-head coil). T1-weighted sMRI scans were acquired through 3D MPRAGE TurboFLASH sagittal sequences with the following characteristics: 2300 ms repetition time; 2.98 ms echo time; 900 ms inversion time; flip angle 9°; 176 slices; 1 mm slice thickness; 256*240 acquisition matrix; 240 Hz/Px bandwidth. sMRI images were segmented using FreeSurfer (v7.1.1) with standard options (recon-all), generating individualised cortical surface meshes of the scalp, cortex, and white matter.

### 2.7 EEG acquisition

High-density EEG data was acquired with a 256-saline-soaked-sponge-sensor, whole-brain Hydrocel Geodesic EGI NetAmp300 system (Electrical Geodesic Inc, Eugene, OR, USA) in an electrically shielded room. Stimulus presentation and event recording synchronisation were managed via E-Prime software (v2.0.8.90, PST, Pittsburgh, PA, USA). Data were sampled at 250Hz, with impedance kept <50KΩ, and Cz as the reference electrode. Electrooculogram (EOG) was recorded from electrodes positioned above, below, and laterally to each eye.

### 2.8 Old/New word recognition task

The stimuli were displayed in white font on a black screen monitor at 1 m viewing distance. Each trial began with a fixation cross followed by a word stimulus presented for 2s (Supplementary Fig. 1). Then a recognition query (“Was this word previously seen?”) was presented, followed by a confidence judgment (“How confident are you in your previous answer?”). The instruction was to answer each question once ready but quickly, using a yes/no button press response without any time limit (once a response was provided, the following step was initiated). Only the responses to the first question (Old/New) were analysed. Four runs of 52 words each (total = 208) with breaks between were presented.

The stimuli were composed of 16 Old words repeated five times (targets, the same words seen in the morning during the FCSRT test), and 64 New words (distractors, not seen before). To characterise the source of EM deficits, in an unbeknownst manner to participants, New words were subdivided into 16 New-Repeated (repeated five times as targets but pertaining generally to semantically unrelated words) and 48 New-Unique words (presented once but pertaining generally to the same semantic categories as targets). The first presentation of a New-Repeated word was coded as New-Unique, leading to 80 Old, 64 New-Repeated, and 64 New-Unique trials. Word presentation order was pseudo-randomised across runs avoiding immediate repetitions (0 to 3 presentations of each repeated word per run) and constant across participants. To minimise repetition bias across sessions, 2 alternating word lists were used (list A for M0/24/48 and B for M12/36/60).

### 2.9 Behavioural data processing

Behavioural data analyses were conducted in R studio (v. 4.2.1), including accuracy, RT, and the main signal detection theory (SDT) metrics. Initial data visualisation suggested potential task-adaptation effects with slower RTs in the first 17 trials of the first run and the first trial of the next three runs; these trials were excluded, in keeping with a balance trial count across the word categories (Old = 76; New-Repeated = 63; New-Unique = 49). The four runs were concatenated, and individual mean RTs of correct responses were computed per experimental condition (Old/New) and session (M0-M60), considering only responses within 150-5000ms post-response-screen-onset. The individual mean accuracy was calculated as the percentage of correct responses per experimental condition and session (*numberofcorrectresponses*/*totalnumberoftrials\**100). SDT metrics included dprime (*d’;* Old/New discrimination sensitivity) and criterion (response bias), as derived from the four SDT response outcomes, i.e., Hits, correct rejections [CR], misses (targets identified as distractors) and false alarms (FAs; distractors identified as targets). We calculated the Hit Rate (*HR* = *Hits*/[*Hits* + *Misses*]) and the FA Rate (*FAR* = *FA*/_[_*FA* + *CR*_]_), transformed it into quantiles of normal distribution (*qnorm*) and we computed *d’* as (*qnorm*[*HR*] − *qnorm*_[_*FA*_]_) and criterion as (−[*qnorm*(*HR*) + *qnorm*(*FA*)]/2).

### 2.10 EEG data preprocessing

EEG data preprocessing was performed with Brainstorm (Version: 3.250506) within MATLAB (v.7.1; Mathworks, Natick, MA, USA).^37^ Previous behavioural findings (see above) led to the exclusion of 12 New and 5 Old trials from the first run and of the first trial from the next 3 runs, to account for task adaptation effects. Preprocessing included removal of flat or noisy channels/periods by visual inspection, 0.1-40Hz bandpass filtering, and trimming of filter-affected time window edges. Epochs were defined around the word stimulus (-200 to 1200ms).^33^ Blinks were automatically detected by Brainstorm using the above-eye electrodes E37 and E18 (or nearby alternatives when these electrodes were not of sufficient quality), followed by a new visual inspection and elimination of problematic periods, channels, and events overlapping with blinks in [-200 to 1200 ms] epochs around word stimuli. Data were then referenced to the average of all channels, reconstructing the signal on Cz. Blink artefacts were corrected using a signal-space projection (SSP) algorithm computed around blink events, typically applying the first projector (blinks) and occasionally the second (saccades). After a final visual inspection to reject any remaining epoch artefacts, events were labelled based on SDT response outcomes (Hits, CR, FA, and Misses, considering responses within 7s maximum after word onset). Finally, for each participant and each session, we averaged the ERPs in response to the correctly recognised Old and New words (i.e., Hits and CR) separately, pooling together the trials from the 4 runs. We also computed overall averages of ERPs across sessions and subjects for figure purposes.

### 2.11 ERP analysis

We performed mean amplitude measurements of ERPs. We defined six time-windows of interest related to known components, based on scientific literature and visual inspection: two reference visual components, P1 (112-116ms) and N1 (152-184ms), and four memory-related components, P3 (252-340ms), FN400 (412-572ms), P600 (620-772ms), and post-retrieval monitoring ERP (872-1040ms) (Friedman and Johnson, 2000; Curran, 2000; Rugg and Curran, 2007; Olichney et al., 2008; Friedman, 2013). Given their distinct topographies, the mean amplitude of P1 and N1 was measured in three regions of interest (ROI), namely, in the occipital lobe (OL) bilaterally and in the parietal lobe (PL). For memory-related components, the ROIs were defined topographically across the following cerebral lobes: bilateral frontal (FL), temporal (TL), and occipital lobe (OL), anterior and posterior PL (Supplementary Fig. 2 and Supplementary Table 1).

### 2.12 Source localisation analysis

Given that individual electrode positions were not collected during data acquisition, we used a standard template of electrode positions for the EEG acquisition system (GSN hydrocel 256 sensor net from EGI), as available in Brainstorm. Electrode positions were co-registered and aligned with the individual sMRI, which were normalised to MNI space using the *spm_maff8* algorithm via a 4×4 affine transformation. Anatomical fiducials (nasion, bilateral preauricular points, anterior and posterior commissures, and inter-hemispheric point) were set as reference to MNI coordinates and used to constrain MRI-MNI alignment. EEG-MRI co-registration involved aligning the template electrode locations on the scalp surface and refining alignment by estimating an optimised rigid-body transformation in order to project electrodes onto the scalp surface. All alignments were visually checked. A Forward Boundary Element Model (BEM, OpenMEEG_BEM v2.4.1) was generated based on head geometry with nested tessellated surfaces of segmented head compartments (1922 vertices per layer; 4mm thickness). With these surfaces, a BEM forward model was created, using the cortex as source space. Noise covariance matrices were computed from pre-stimulus baseline segments (-200 to 0 ms, average subtracted) across all epochs for each subject, session, and run separately. Sources with unconstrained orientations were computed using weighted minimum norm estimation (wMNE), in the form of cortical current density maps (pA.m) and averaged across trials and runs in each condition of interest (Hits and CR), for each subject and session (M0-M60). They were standardised by z-transforming cortical current time series relative to the pre-stimulus baseline activity, and the norm of the unconstrained sources was computed at each vertex. For group-level analysis, these individual maps were averaged per group (controls, stable, progressors) and session. For statistical analyses, the mean source amplitudes were extracted from 22 predefined cortical regions of the Automated Anatomical Labeling (AAL) atlas (Supplementary Fig. 3), selected for their relevance to recognition memory processes and AD.^40^ This extraction was performed in each time-window of interest (252-340ms [P3], 412-572ms [FN400], 620-772ms [P600], and 872-1040ms [post-retrieval monitoring]) for each subject, session, and condition (Hits / CR).

### 2.13 Statistical analyses

All statistical analyses were conducted in R Studio (v. 4.4.3; R Core Team, 2025).

#### 2.13.1 Demographic and clinical assessments at baseline (M0)

Baseline group differences in age, education level, and self-reported SMCs were assessed using one-way Welch ANOVAs (‘car’ R package, v.3.1.0), with group (controls, stables, and progressors) as fixed between-subject factor. Sex ratio was compared across the 3 groups using a chi-square test. We further assess Aβ_SUVr differences across the Aβ+ groups (stables vs. progressors) and FDG_SUVr differences across the N+ groups (stables_Aβ+_N+ vs. progressors Aβ+_N+) using t-test. Baseline inclusion neuropsychological scores (MMSE, FAB, FCSRT_TR, McNair) were evaluated using general linear models (glm; ‘stats’ R package, v.4.2.1) with group as fixed between-subject factor. Age at baseline (M0) was included as a mean centred numeric covariate and education level and sex were included as categorical covariates, to account for potential confounds. Statistical significance was set at p<0.05 with multiple comparison correction, and Games-Howell post-hoc tests (‘rstatix’ R package, v.0.7.0) were used where appropriate.

#### 2.13.2 Behavioural performance in the Old / New word recognition task

We used linear mixed-effects models (LMMs; lme4, v.1.1.29), which allow accounting for repeated measurements and missing/unbalanced data, to assess the longitudinal behavioural outcomes (accuracy, *d’*, criterion, and log-transformed RT) in the word recognition task. For accuracy, a generalised version of LMM (GLMM) with a binomial logistic link was used to account for the binary structure of the data. For accuracy and (log-transformed) RT, the models included the following fixed-effect factors: word-category (Old/New), session (M0-M60) and their interaction as within-subject factors; group (controls, stables, progressors) and its interactions with word-category and session as between-subject factors; age, sex, and education as additional covariates of no interest. The random-effect structure comprised random intercept and random effects of the word-category and session factors across participants. Similar LMMs were used for *d’* and criterion, but without word-category as factor and with only random intercept across participants as random-effect structure (due to limited observations). Statistical significance was reported in the form of p values, derived from chi-square likelihood-ratio tests for accuracy analysis (GLMM) and from F-tests with Satterthwaite approximations for the remainder analysis (LMMs). Post-hoc comparisons for significant within- or between-factor fixed effects and interactions were performed using emmeans (R package, v.1.7.5), using multivariate t-distribution corrections to control the family-wise error rate for multiple comparisons (*α* = 0.05) (Tzourio-Mazoyer et al., 2002).

#### 2.13.3 ERP sensor-level analysis of the Old/New word recognition task

We used LMMs to assess the mean amplitude differences in the predefined ERP time-windows—P300 (252–340 ms), FN400 (412–572 ms), P600 (620–772 ms), and post-retrieval monitoring (872–1,040 ms)—extracted from eight ROIs (bilateral FL, TL, and OL; and anterior and posterior PL). For each time-window and ROI, we analysed amplitude differences across experimental conditions and groups using LMMs with word-category, session and their interaction as within-subject fixed-effect factors, group and its interactions with word category and session as between-subject factors, age, sex and education as additional fixed-effect covariates. Random intercept and random effects of word category and session across participants were also included. Significant findings were followed by post-hoc pairwise comparisons using the emmeans function with multivariate t-distribution correction for multiple comparisons. In addition, we applied false discovery rate (FDR) correction to adjust for multiple comparisons across ROIs in each ERP time-window.

#### 2.13.4 ERP source-level analysis of the Old / New word recognition task

Source-reconstructed activities were evaluated using the same set of time-windows and statistical framework as in the ERP analyses. Mean source amplitudes were extracted from the 22 predefined cortical regions from the AAL atlas. For each region and time-window, LMMs were fitted with word-category and session as within-subject fixed-effects, group as a between-subject fixed-effect factor, also including the interactions between these factors, and age, sex, and education as additional fixed-effect covariates. The same random-effect structure and post-hoc pairwise comparisons using emmeans and corrections for multiple comparisons as detailed above were performed when significant fixed-effects effects or interactions were encountered.

## 3. Results

### 3.1 Demographic and neuropsychological characteristics of the groups and clinical relation with biomarkers

From the INSIGHT-preAD cohort, we considered progressors to prodromal AD (15 progressors), and matched them in age, sex and education level at M0 to a group of 15 stable participants and a group of 15 controls. This subsample had an overall mean age of 77.4 ± 3.3 years [76-79 years] at baseline, was predominantly female (56% [*n*=25]) and highly educated (*mean NSC*: 6.4 ± 2.1, i.e., undergraduate level; no *NSC*=1; Table 1). The stable group was matched as much as possible to the progressors in Aβ- and FDG-SUVr. The Aβ-SUVr significantly differed between the Aβ+ groups [*t*(26.2)=2.97, *p*=0.006], with higher values in the progressors than the stable participants (Table 1). Neurodegeneration (FDG-SUVr) did not differ significantly between groups. Furthermore, no significant group difference was found at baseline in self-reported SMC at consultation, although controls reported the highest average scores. All groups performed within normal age-adjusted ranges on neuropsychological total scores: *MMSE*=28.4±0.8 (≥27/30); the *FAB*=16.4±1.7 (≥16/18), and *FCSRT-TR*=45.7 ± 2.2 (≥41/48). Due to technical issues, diagnosis, death, or study withdrawal, the sample size declined during follow-up, which at M60 resulted in 14 controls, 12 stable Aβ+, and 8 progressor participants (Supplementary Table 2).

**Table 1.**
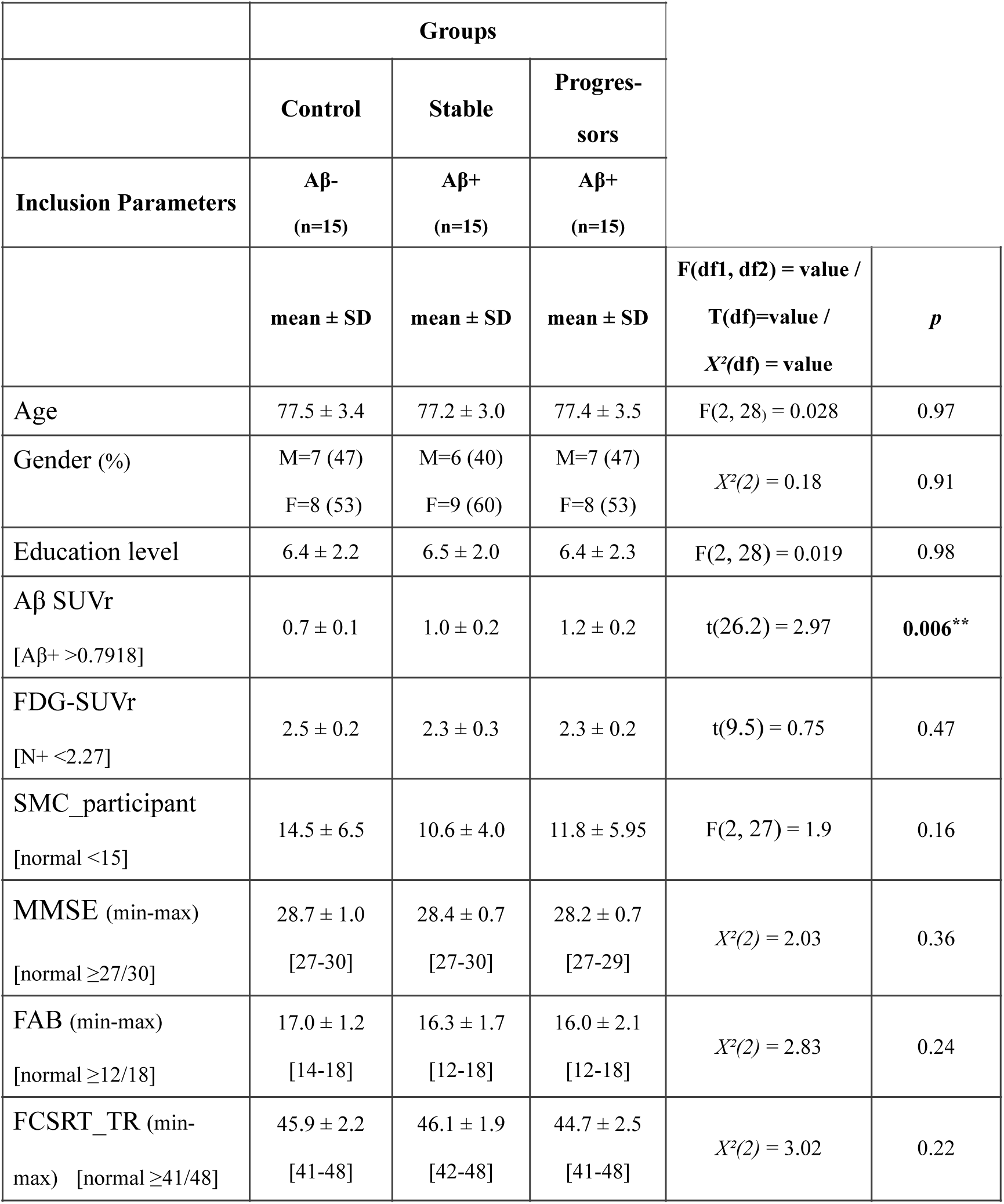
Demographic and clinical characteristics of the control, stable, and progressor groups at baseline (M0).

|  | Groups |  |  |  |  |
| --- | --- | --- | --- | --- | --- |
|  | Control | Stable | Progressors |  |  |
| Inclusion Parameters | Aβ-<br>(n=15) | Aβ+<br>(n=15) | Aβ+<br>(n=15) | | |
| | mean ± SD | mean ± SD | mean ± SD | F(df1, df2) = value /<br>T(df)=value /<br>X<sup>2</sup>(df) = value | *p* |
| Age | 77.5 ± 3.4 | 77.2 ± 3.0 | 77.4 ± 3.5 | F(2, 28) = 0.028 | 0.97 |
| Gender (%) | M=7 (47)<br>F=8 (53) | M=6 (40)<br>F=9 (60) | M=7 (47)<br>F=8 (53) | X<sup>2</sup>(2) = 0.18 | 0.91 |
| Education level | 6.4 ± 2.2 | 6.5 ± 2.0 | 6.4 ± 2.3 | F(2, 28) = 0.019 | 0.98 |
| Aβ SUVr<br>[Aβ+ >0.7918] | 0.7 ± 0.1 | 1.0 ± 0.2 | 1.2 ± 0.2 | t(26.2) = 2.97 | **0.006\*\*** |
| FDG-SUVr<br>[N+ <2.27] | 2.5 ± 0.2 | 2.3 ± 0.3 | 2.3 ± 0.2 | t(9.5) = 0.75 | 0.47 |
| SMC_participant<br>[normal <15] | 14.5 ± 6.5 | 10.6 ± 4.0 | 11.8 ± 5.95 | F(2, 27) = 1.9 | 0.16 |
| MMSE (min-max)<br>[normal ≥27/30] | 28.7 ± 1.0<br>[27-30] | 28.4 ± 0.7<br>[27-30] | 28.2 ± 0.7<br>[27-29] | X<sup>2</sup>(2) = 2.03 | 0.36 |
| FAB (min-max)<br>[normal ≥12/18] | 17.0 ± 1.2<br>[14-18] | 16.3 ± 1.7<br>[12-18] | 16.0 ± 2.1<br>[12-18] | X<sup>2</sup>(2) = 2.83 | 0.24 |
| FCSRT_TR (min-max)<br>[normal ≥41/48] | 45.9 ± 2.2<br>[41-48] | 46.1 ± 1.9<br>[42-48] | 44.7 ± 2.5<br>[41-48] | X<sup>2</sup>(2) = 3.02 | 0.22 |

The values of the main inclusion scores are presented as mean ± SD or number (and % in parentheses). P-values reflect the main effect of group from Welch ANOVA, chi-square test, or general linear models (GLMs) performed for each variable as detailed in Methods. Abbreviations: SD = Standard Deviation; Aβ SUVr = β-Amyloid Standardized Uptake Value ratio; df = degrees of freedom; MMSE = Mini Mental State Examination; FAB = Frontal Assessment Battery; FCSRT = Free and Cued Selective Reminding Test. The p value of the statistical analysis performed between the 3-groups at session M0 is reported for each outcome variable with ***: p < 0.001; **: p≤0.01; *: p≤0.05.

### 3.2 Behavioural performance

Behavioural performance in the Old/New word recognition task was assessed using accuracy, *d’* and criterion indices of SDT, and RT. As we already observed previously in an analysis of the whole cohort (Raposo-Pereira et al., 2025), accuracy remained near ceiling across sessions in controls (*M0* = 0.97 ± 0.03; *M60* = 0.98 ± 0.02) and stable participants (*M0* = 0.98 ± 0.02; *M60* = 0.97 ± 0.02), while it declined notably in progressors (*M0* = 0.93 ± 0.10; *M60* = 0.78 ± 0.17; Fig. 1A). There was a significant interaction between word-category and session (*χ²*(5) = 17.60, *p* = 0.003) accompanied by significant interactions between word-category and group (*χ²*(2) = 10.05, *p* = 0.007), session and group (*χ²*(10) = 72.95, *p* < 0.0001), and a 3-way interaction between group, word-category, and session (*χ²*(10) = 72.21, *p* < 0.0001). This reflected a lower performance in the progressors than controls at recognising New words in all sessions except M36 and Old words only in sessions M48 and M60. When compared with the stable participants, the progressors showed lower performance at recognising New words at sessions M48 and M60, and Old words at sessions M12, M48, and M60. Accuracy varied with education level (*χ²*(6) = 17.65, *p* = 0.007) and age (*χ²*(1) = 7.92, *p* = 0.005).

**Figure 1.**
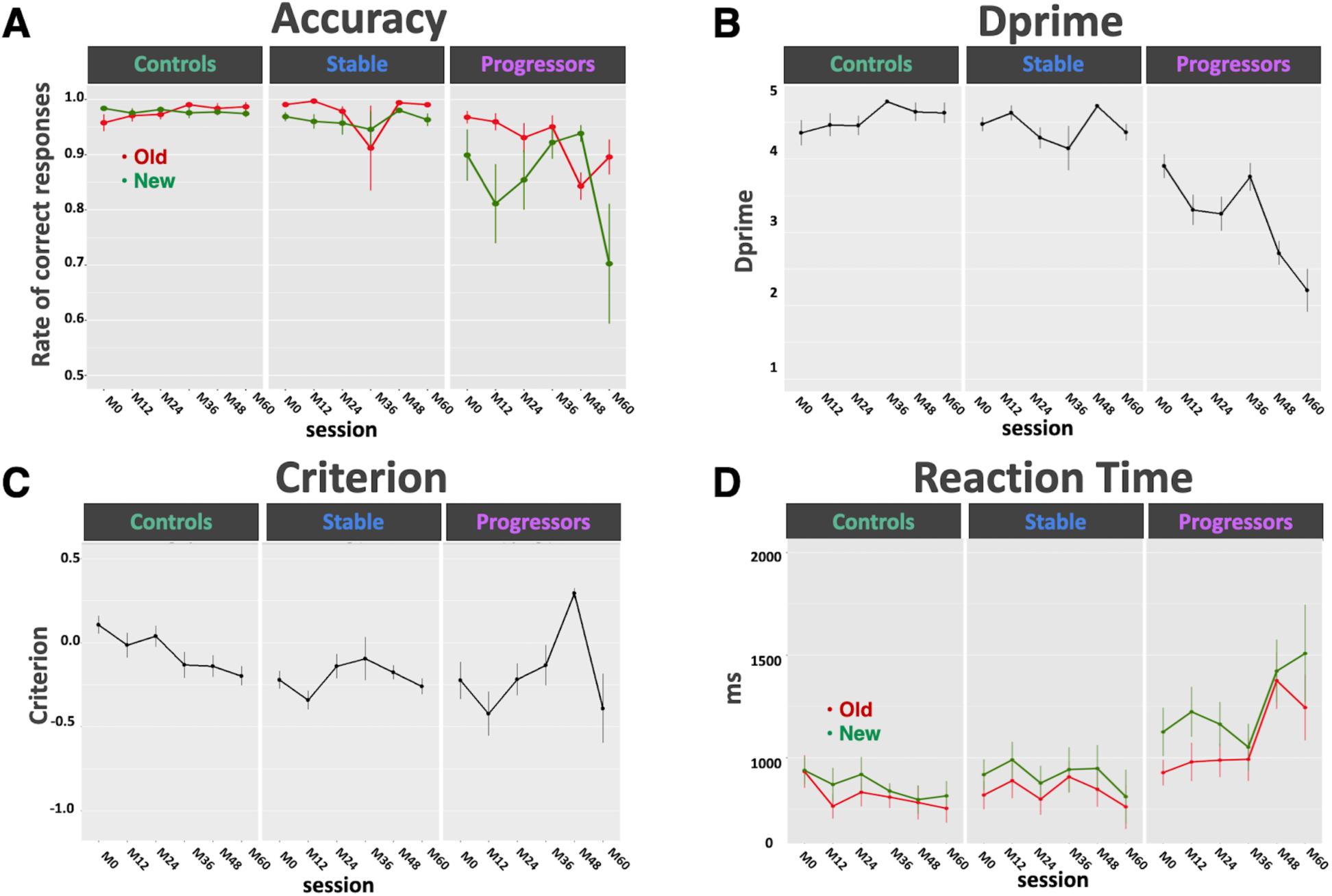
Behavioural performance (accuracy, dprime [*d’*], criterion, and reaction time [RT]) in the Old/New word recognition task across sessions for each group. Each plot represents the mean values for each session (M0-M60), for each group (controls, stable, and progressors), and for each condition (Old and New) in the case of accuracy and RT; the vertical lines represent the standard errors of the mean. (**A)** accuracy (rate of correct responses). (B) *d’* (Old/New discrimination sensitivity). (C) criterion (response bias). **(D)** RT (ms). Old words are in red and New words are in green.

The analysis of *d’* (Old/New word discrimination sensitivity) showed a main effect of group (*F*(2,33) = 21.98, *p* < 0.0001), with progressors performing significantly worse than both controls and stable participants (both p < 0.0001; Fig. 1B). There was a significant effect of session (*F*(5,416) = 8.18, *p* < 0.0001) and an interaction between session and group (*F*(10,417) = 11.30, *p* < 0.0001). Post-hoc comparisons indicated reduced *d’* in progressors vs. controls from M12 onward, and in progressors vs. stable participants at M12, M24, M48, and M60. This echoed the steeper decline of accuracy across sessions in the progressors relative to both other groups.

Criterion analysis showed a main effect of session (*F*(5,416) = 5.74, *p* < 0.0001) and an interaction between session and group (*F*(10,416) = 3.34, *p* = 0.003; Fig. 1C). Overall, all groups exhibited a negative response criterion, indicating a bias toward “yes” (“old”) responses, resulting in higher hit rates but also increased false alarms. The controls were the only group with a positive baseline criterion, which then decreased significantly, becoming negative, from session M0 to M60 (*p* = 0.03). The stable group maintained a consistently negative criterion longitudinally. In contrast, the progressors showed pronounced longitudinal variability: first a negative criterion that significantly increased towards positivity from M12 to M48 (*p* = 0.0001), followed by an abrupt decline to a negative criterion from M48 to M60 (*p* < 0.0001).

Mean RTs were longer in progressors from session M0 (1039 ± 342 ms) as compared to controls (935 ± 266 ms) and stable participants (879 ± 275 ms; Fig. 1D). RT analysis revealed a main effect of group (*F*(2,38) = 8.27, *p* = 0.001) and post-hoc tests showed significantly longer RTs in progressors relative to controls (*p* = 0.0009). There was also a main effect of word-category (*F*(1,35) = 26.33, *p* < 0.0001), with faster responses for Old than New words. An interaction between session and group (*F*(10,38) = 4.28, *p* = 0.0005) indicated that the progressors were significantly slower than the controls from session M36 onwards and significantly slower than the stable participants at M60. Additionally, RTs increased with age (*F*(1,81) = 4.43, *p* = 0.04), were longer in males (*F*(1,32) = 10.25, *p* = 0.003), and varied with education (*F*(6,29) = 2.7, *p* = 0.03). No other significant difference was found.

### 3.3. Sensor-level analysis of ERPs

As can be seen in Fig. 2A, 2B and 2C, the grand mean of the ERPs across subjects and conditions allowed visualising the typical ERP components related to the processing of words, memory, and the Old/New effect.

**Figure 2.**
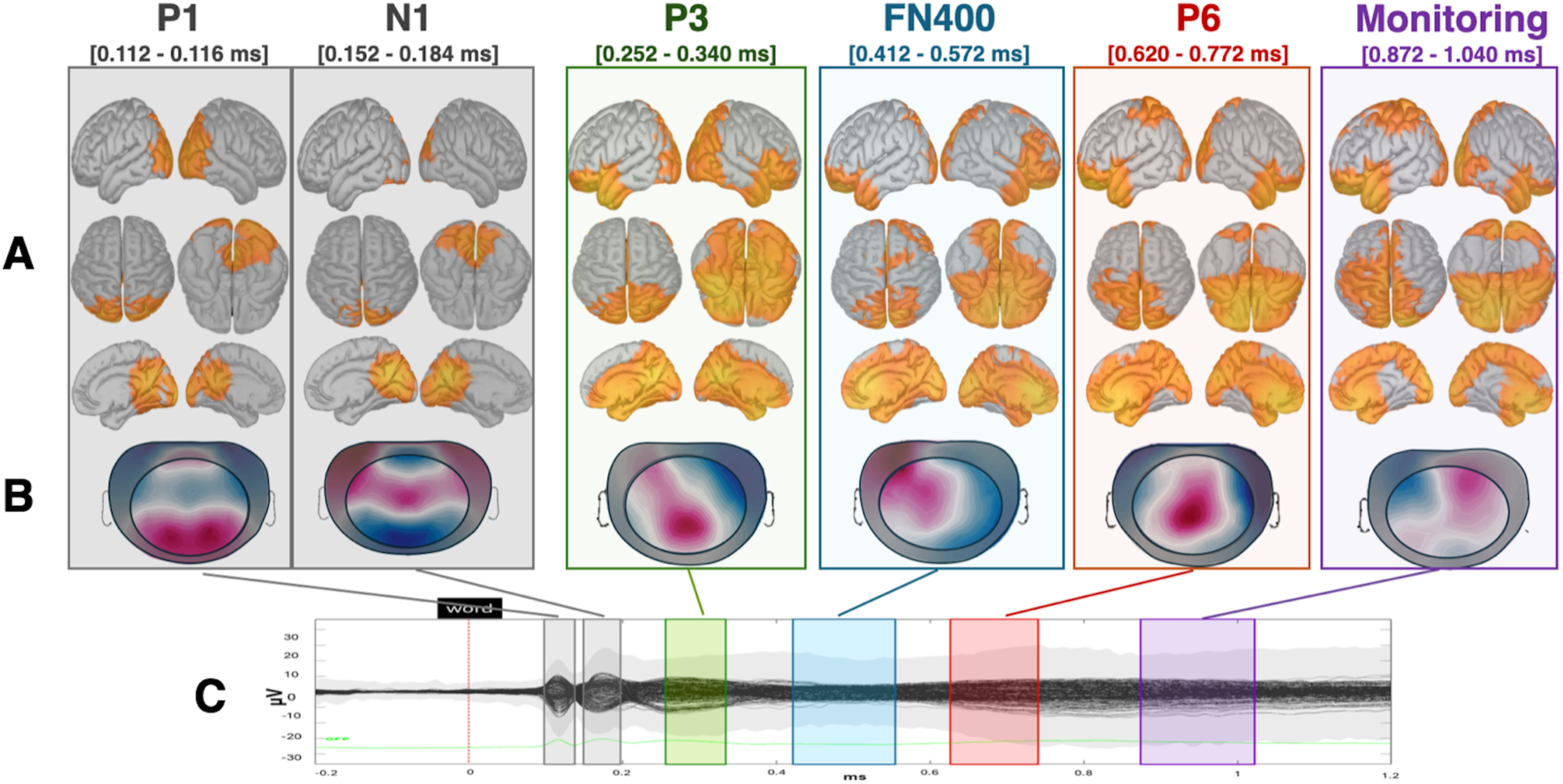
Cortical source activity and scalp topographical maps of event-related potential (ERP) components across participants, word-category (Old/New), and sessions (M0-M60) during the Old/New word recognition task. **(A)** Source-level cortical activation maps shown in left and right sagittal views (first row), dorsal and ventral views (second row), and right and left medial views (third row) for each ERP component. **(B)** Mean scalp topographical maps for each ERP component; red indicates positive potentials and blue negative potentials. The black circle outlines the electrode array assessed. **(C)** Grand-average ERP time course on all overlaid electrodes; the ERPs time-locked to word onset (0 ms) were averaged across all participants, sessions, and word categories. Color-shaded windows denote the latency ranges of the ERP components of interest: two reference visual components, P1 (112-116ms) and N1 (152-184ms), and four memory-related components, P3 (252–340 ms), FN400 (412–572 ms), P600 (620–772 ms), and post-retrieval monitoring (872–1040 ms).

LMMs were fitted separately on the mean amplitude of ERPs on 8 ROIs covering the scalp (bilateral FL, TL, OL, and anterior and posterior PL) in the time-windows of the memory-related ERP components: 252–340 ms (P300), 412–572 ms (FN400), 620–772 ms (P600), and 872–1040 ms (post-retrieval monitoring). We observed a significant main effect of word-category across several ROIs and time-windows in line with the canonical Old/New ERP effect. In line with the attended topographical distribution of familiarity (FN400), recollection (P600), and monitoring-related processes (Fig. 3 and Supplementary Table 3), Old words elicited greater amplitudes than New words, predominantly in left and central-posterior regions, whereas New words elicited greater amplitudes than Old words predominantly in the right FL (Fig. 3 and Supplementary Table 3).

**Figure 3.**
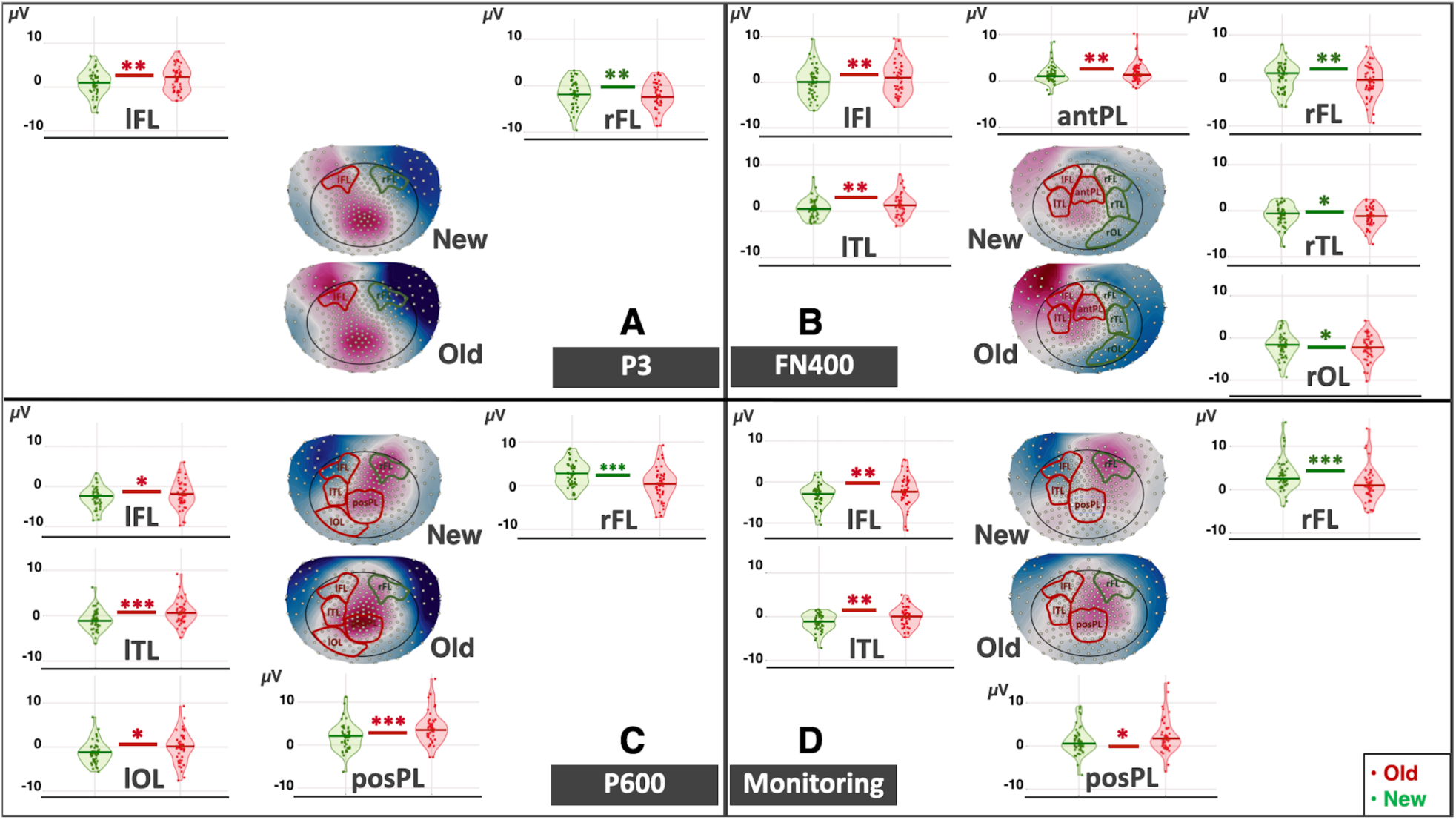
Topographical maps and regional effects during the Old/New word recognition task for each event-related potential (ERP) component of interest. Panels represent each ERP component: **(A)** P3 **, (B)** FN400**, (C)** P600**, (D)** post-retrieval monitoring. Violin plots display mean ERP amplitudes (*µV*) in regions of interest where significant Old/New differences were found (Old, red; New, green). Dots represent the participant-level mean amplitude across sessions (M0-M60). Significant differences are denoted by asterisks (*= *p* <.05, ** =*p* <.01, ***=*p* <.001), with colour indicating the direction of the Old-New effect (red: Old>New; green: New>Old). The topographical maps show grand-averages for each condition (New on top, Old on bottom), with outlined regions of interest: left Frontal Lobe (lFL); right Frontal Lobe (rFL); anterior Parietal Lobe (antPL); posterior Parietal Lobe (posPL); left Temporal Lobe (lTL); right Temporal Lobe (rTL); left Occipital Lobe (lOL); right Occipital Lobe (rOL).

Only one ROI—the left OL—showed significant group-related effects across ERP time-windows (Fig. 4). An interaction between word-category and group emerged in the FN400 (*F*(2,13)= 5.39, *p* = 0.02), P600 (*F*(2,15) = 8.33, *p* = 0.004), and post-retrieval monitoring (*F*(2,19) = 4.97, *p* = 0.02) time-windows. However, after correcting for multiple comparisons only the P600 effect remained significant. Post-hoc analyses revealed that while the controls and stable participants showed the expected Old>New amplitude pattern overall, this pattern was attenuated and inverted in the progressors at session M60. Additionally, a three-way interaction between word category, group, and session was observed in the FN400 (*F*(10,15) = 2.97, *p* = 0.03), P600 (*F*(10,19) = 3.55, *p* = 0.009) and post-retrieval monitoring (*F*(10,26) = 3.23, *p* = 0.008) time-windows. Despite not surviving multiple comparisons, post-hoc tests showed this three-way interaction was driven by the progressors, at M60, with a marked New>Old amplitude pattern (all *p* < 0.0001).

**Figure 4.**
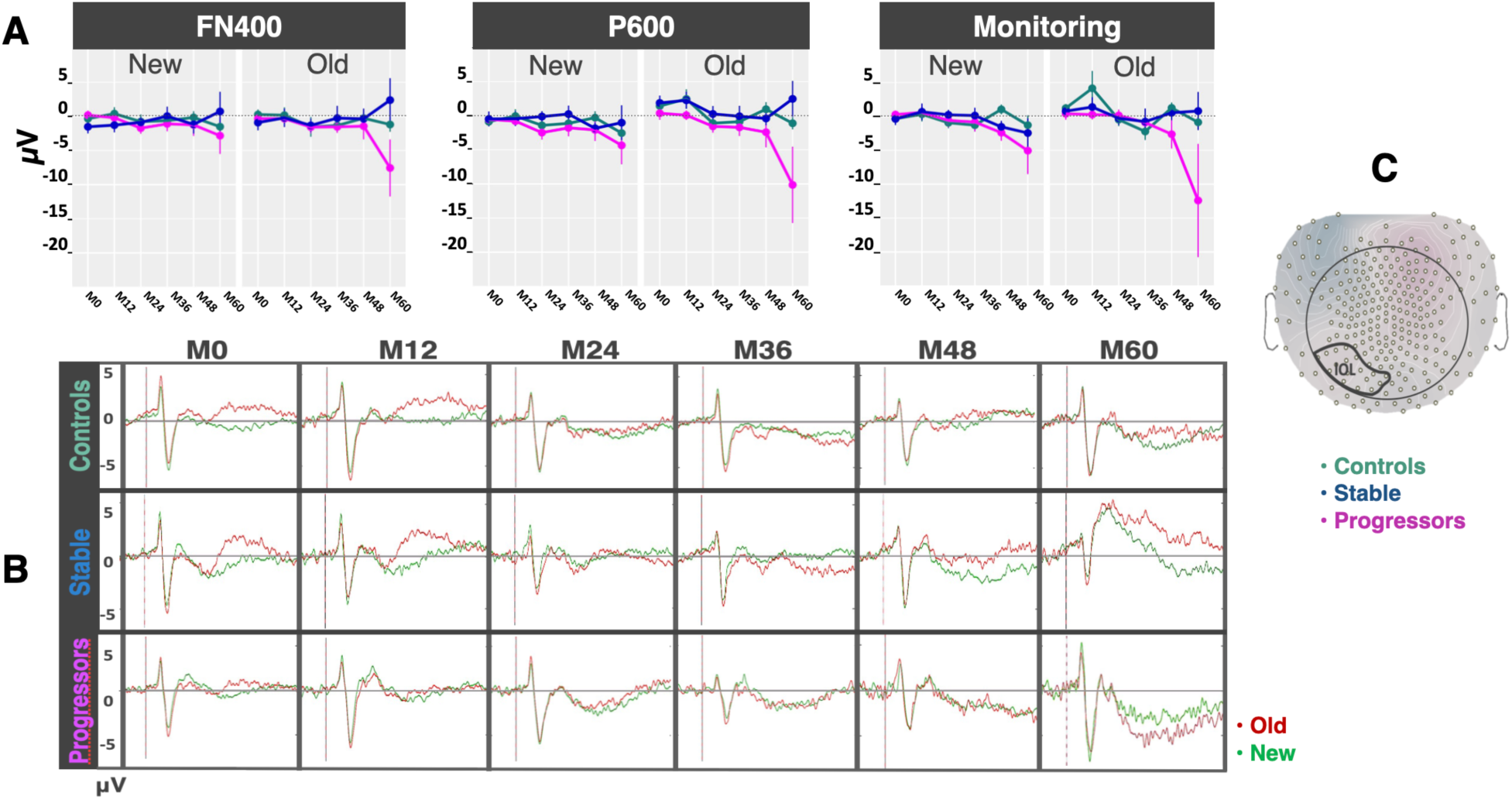
Longitudinal group differences in event-related potentials (ERP) waveforms, and mean amplitudes across word-categories and components in the left Occipital Lobe (lOL) during an Old/New word recognition memory task. **(A)** Mean amplitudes (*µV*) for the FN400, P600, and post-retrieval monitoring components during Old and New word-category across sessions (M0 to M60), across groups: Controls (turquoise), Stable (blue), and Progressors (magenta). **(B)** Grand-average ERP waveforms (Old: red; New: green) for each group and session (rows: groups, columns: sessions). Progressors show a progressive decline in the recognition of Old words especially in the P600 and post-retrieval monitoring components when compared to the Controls and Stable. **(C)** The topographical map indicates the lOL region where significant differences between the groups were observed.

### 3.4 Source-level analysis of ERPs

At the cortical source level, there was a significant main effect of group in the left inferior FL in the four time-windows of interest (P300: [*F*(2,36)= 4.77, *p* = 0.015]; FN400: [*F*(2,36)= 6.24, *p* = 0.005]; P600: [*F*(2,28)= 5.73, *p* = 0.008]; post-retrieval monitoring: [*F*(2,28)= 4.11, *p* = 0.027]; Fig. 5), as well as in the right Posterior Cingulate Cortex (PCC; P300: [*F*(2,37)= 3.44, *p* = 0.043]; FN400: [*F*(2,32)= 6.94, *p* = 0.003]; P600: [*F*(2,32)= 6.60, *p* = 0.004]; post-retrieval monitoring: [*F*(2,30)= 6.35, *p* = 0.005]; Fig. 5). Post-hoc analysis consistently revealed reduced activations in these areas in the progressors as compared to the controls (left inferior FL: P300, *p* = 0.02; FN400, *p* = 0.004; P600, *p* = 0.006; post-retrieval monitoring: *p* = 0.02; PCC: P300, *p* = 0.05; FN400, *p* = 0.003; P600, *p* = 0.004; post-retrieval monitoring: *p* = 0.005).

**Figure 5.**
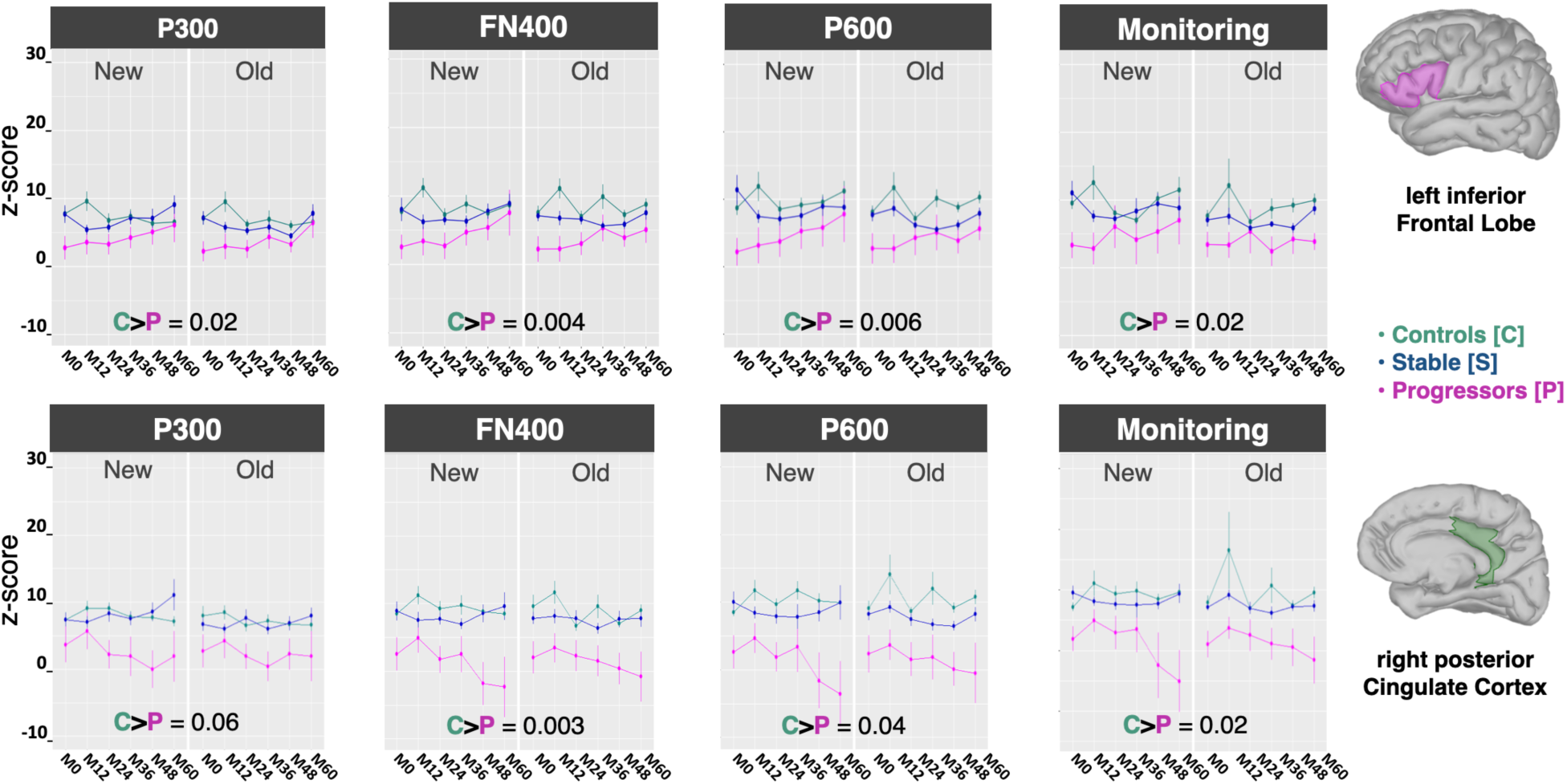
Time-course of source activity in regions showing significant group differences across all memory-related event-related potential (ERP) time windows. Mean source amplitudes (z-score) are plotted across four ERP time-windows (P300, FN400, P600, and post-retrieval monitoring) for Old and New words across sessions (M0–M60), in the left inferior frontal lobe (FL; top row) and right posterior cingulate cortex (PCC; bottom row). Controls (C; in turquoise) consistently showed greater activation than Progressors (P; in magenta) with the most pronounced and sustained differences observed in the P600 window. Significant post-hoc group differences are reported within each plot. Stable Aβ+ participants (in blue) showed intermediate activation levels. These patterns suggest progressive disruption of recollection and monitoring-related processes in progressors, consistent with early network failure in AD-vulnerable regions.

In the P600 time-window, significant hypoactivations were further observed in the progressors versus the controls in the left inferior and superior TL regions ([*F*(2,31)=4.17, *p*=0.02] and [*F*(2,31)=3.91, *p*=0.03], respectively), right medial TL (*F*(2,26)=4.98, *p*=0.015), right PCC (*F*(2,32)=6.60, *p*=0.004), and left and right precuneus ([*F*(2,27)=4.29, *p*=0.02] and [*F*(2,28)=3.93, *p*=0.03], respectively; Fig. 6).

**Figure 6.**
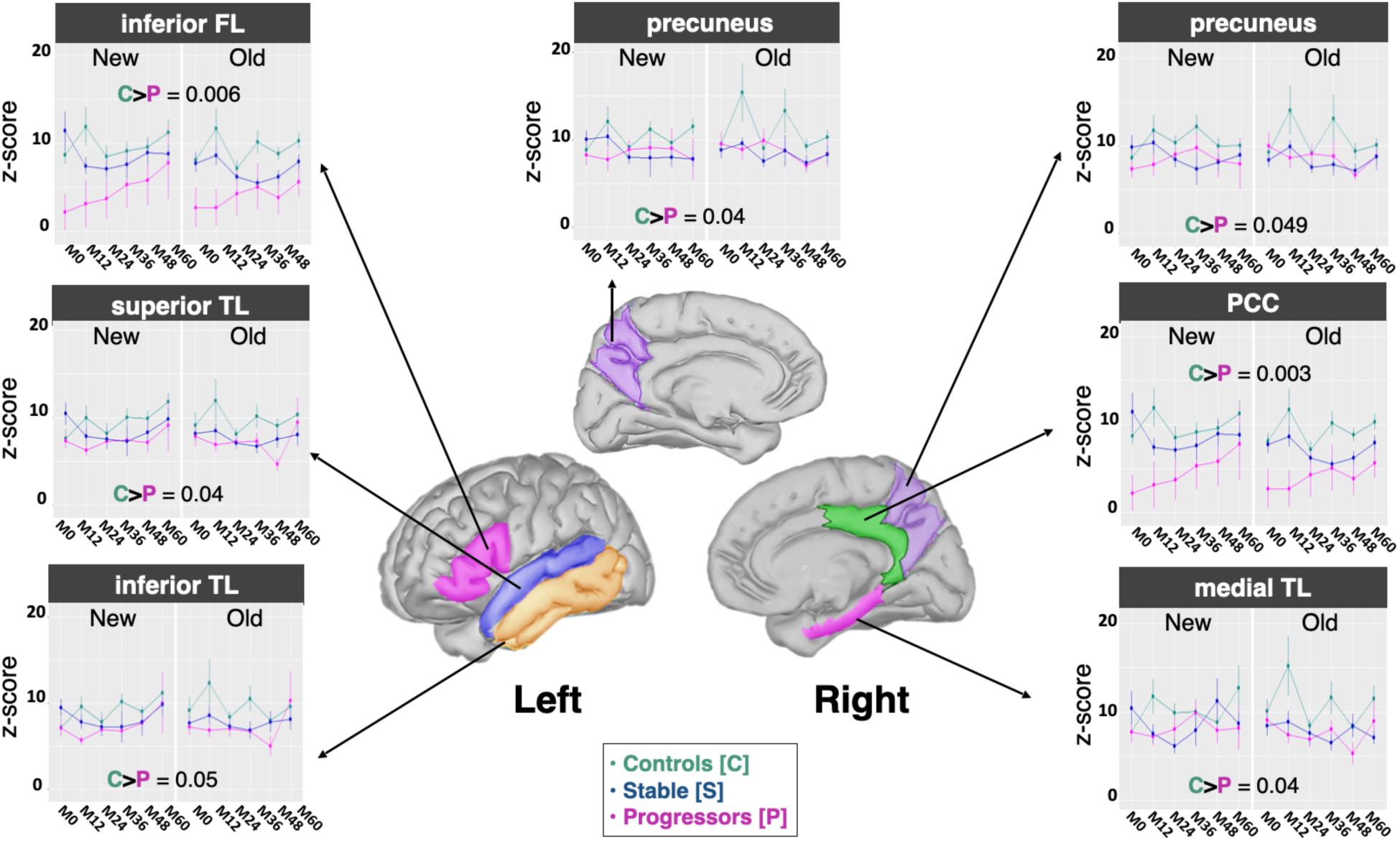
Source activation differences between controls and progressors in the P600 time window. The plots show the mean amplitude of sources in different regions in the P600 time window for the three groups. The progressors (P; in magenta) showed significantly reduced source activity compared with controls (C; in turquoise) in the left inferior frontal lobe (FL), left superior and inferior temporal lobe (TL), bilateral precuneus, right posterior cingulate cortex (PCC), and right medial TL. Line graphs show mean source amplitude (in z-scores) across sessions (M0 to M60), for each group (controls, stable [blue], and progressors), for New and Old words respectively. Error bars indicate standard error of the mean across subjects. Significant differences between groups in post-hoc comparisons are reported within each plot. The middle brain views show the anatomical locations of the regions displayed in the line plots.

Moreover, significant hypoactivations in the stable group relative to the controls were also observed exclusively in the P600 time-window in the left rostral-middle FL (*F*(2,47)=4.85, *p*=0.01), left OL(*F*(2,37)=4.18, *p*=0.02), right inferior TL (find above), and left and right anterior cingulate cortex (ACC; [*F*(2,31)=4.94, *p*=0.01] and [*F*(2,27)=4.09, *p*=0.03], respectively; Fig. 7).

**Figure 7.**
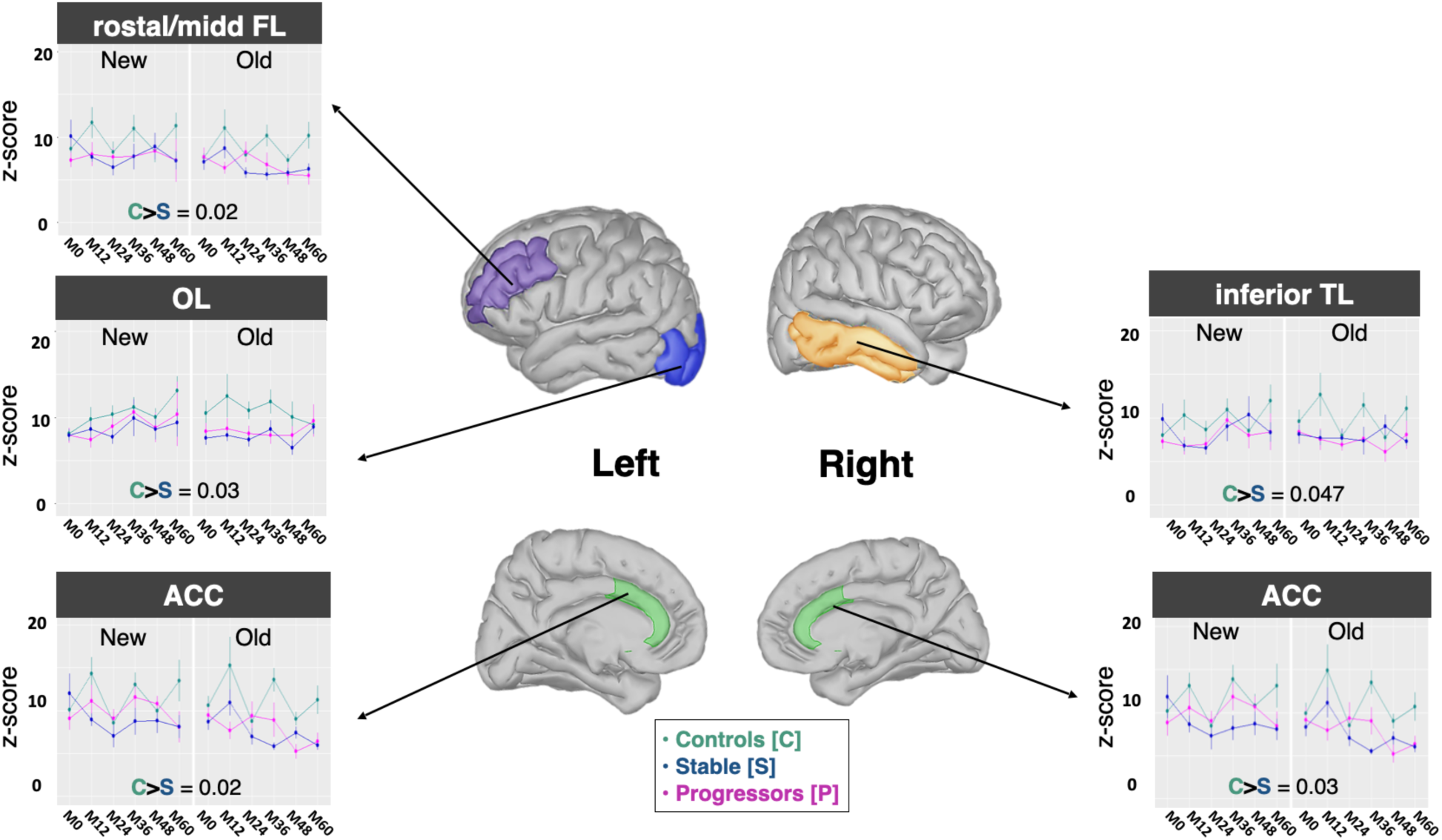
Source activation differences between controls and stable (Aβ+) individuals in the P600 time window. The plots show the mean amplitude of sources in different regions in the P600 time window for the three groups. The Stable Aβ+ participants (S; in blue) showed significantly reduced source activity as compared with the Controls (C; in turquoise) in the left rostral/middle Frontal Lobe (rostral/midd FL), left occipital lobe (OL), bilateral anterior cingulate cortex (ACC), and right inferior temporal lobe (TL). Line plots display mean source amplitudes (in z-scores) across sessions (M0 to M60), for each group (controls, stable, and progressors [magenta]), for New and Old words. Error bars indicate standard error of the mean across subjects. Significant differences between groups in post-hoc comparisons are reported within each plot. The middle brain views show the anatomical locations of the regions displayed in the line plots.

## 4. Discussion

In this retrospective analysis, the INSIGHT pre-AD cohort participants who later progressed to AD, despite being cognitively normal at baseline, showed subtle behavioural and electrophysiological abnormalities during an Old/New word recognition task, compared to stable Aβ+ participants and healthy controls.

Behaviourally, progressors showed slower RTs alongside lower accuracy and *d’* scores, suggesting early deficits in retrieval and decision-making and diminished recognition specificity (Grober et al., 2000; Friedman and Johnson, 2000; Dubois et al., 2018; Raposo Pereira et al., 2024b). Slower responses likely reflected increased cognitive effort and reduced processing efficiency, while lower discrimination performance suggested weakened encoding or retrieval accuracy, biasing progressors towards familiarity-based judgments and reliance on guessing responses (Grober et al., 2000; Friedman and Johnson, 2000; Dubois et al., 2018; Raposo Pereira et al., 2024b). These subtle behavioural declines in memory precision and processing speed, align with evidence that early synaptic and cognitive dysfunction emerge years before AD diagnosis (Sperling et al., 2011; Dubois et al., 2018; Busche and Hyman, 2020; Hampel et al., 2021).

At the sensor-level, progressors exhibited reduced FN400, P600, and post-retrieval monitoring amplitudes, indexing disruption across distinct retrieval-related subprocesses. Attenuated FN400 amplitudes suggest impaired context-free familiarity-based processing and semantic integration, whereas attenuated P600 amplitudes suggest compromised recollection, consistent with inefficient retrieval of contextual details (Friedman, 2013; Olichney et al., 2011; Musaeus et al., 2019). These results are consistent with prior research, showing that FN400 and P600 effects are sensitive markers of early AD-related memory dysfunction, in particular, an attenuated P600 to word repetition, likely reflecting impaired recollection and memory updating that predicts progression from MCI to AD (Friedman and Johnson, 2000; Rugg and Curran, 2007; Olichney et al., 2008; Horváth et al., 2018; Paitel et al., 2021; Jessen et al., 2020; Wang et al., 2024). Diminished late monitoring activity further denoted weakened evaluative and control processes during recognition decisions (Roper, 2011; Razafimahatratra et al., 2023). Within task context, these alterations likely reflect degraded episodic memory (EM) representations and impaired top-down modulation, leading to incomplete reactivation of perceptual and contextual details required for accurate old/new discrimination and confidence judgments (Olichney et al., 2011; Olichney et al., 2013; Paitel et al., 2021; Razafimahatratra et al., 2023). Notably, group differences were restricted to the left parieto-occipital region, a hub for integrating visual-perceptual input with memory representations (Friedman, 2013; Tsivilis et al., 2015; Musaeus et al., 2019). Since posterior cortices are often affected early in AD, these findings suggest accelerated disruption of bottom-up sensory processing and compromise recognition memory performance (Braak and Braak, 1991; Jack et al., 2010; Palmqvist et al., 2017; Busche and Hyman, 2020; Hampel et al., 2021; Zamani et al., 2025). No other significant electrophysiological differences between the groups were found.

Source-level analyses revealed widespread hypoactivation across nodes of the EM network and default mode network (DMN) in the progressors. Compared with controls, reduced activity was observed in the left IFG and right PCC across all memory-related ERP time windows, suggesting deficits in strategic retrieval, semantic memory, and episodic processing (Sperling et al., 2011; Palmqvist et al., 2017; Musaeus et al., 2019; Wang et al., 2024). During recollection-specific processing (P600), hypoactivation extended to the left TL, right MTL and bilateral precuneus consistent with disrupted hippocampal-cortical interactions critical in semantic and retrieval efficiency (Olichney et al., 2011; Tsivilis et al., 2015; Musaeus et al., 2019). These patterns align with evidence of early disruptions in DMN and semantic networks observed in mild cognitive impairment (MCI) individuals and preclinical AD during memory retrieval (Rugg and Curran, 2007; Sperling et al., 2011; Dubois et al., 2018; Musaeus et al., 2019; Wang et al., 2024; Liang et al., 2025).

Age-related left-to-right posterior shifts in ERP old/new effects have been interpreted as compen^-^ satory recruitment in response to degeneration of the typically left-lateralised recollection processes, consistent with electrophysiological and MRI studies in healthy aging and early AD (Rugg and Curran, 2007; Musaeus et al., 2019; Cabeza et al., 2018; Zamani et al., 2025). The mismatch between scalp and source-level activity may likewise indicate transiently compensatory anterior mechanisms, that ultimately fails as typical AD posterior network dysfunction progresses (Olichney et al., 2011; Busche and Hyman, 2020; Musaeus et al., 2019; Paitel et al., 2021; Zamani et al., 2025). These mechanisms would be differentially captured at sensor versus source levels. The origin of this discrepancy is unclear but it may stem from normalisation steps of source localisation, which introduce a non linearity between sensor-level and source-level, reconstructed, activities, with different signal-to-noise ratio or noise sensitivity at each level.

In contrast, stable Aβ+ participants who remained cognitively normal, showed only mild localised P600 hypoactivations in regions related to cognitive control and perceptual-semantic integration—left rostral FL and OL, right inferior TL; and bilateral ACC (Musaeus et al., 2019; Liang et al., 2025). This might reflect early subclinical inefficiencies while preserving broader network function, consistent with neural resilience and compensatory capacity as moderators of AD progression (Sperling et al., 2011; Stern, 2012; Dubois et al., 2018; Cabeza et al., 2018; Arenaza-Urquijo and Vemuri, 2018; Hampel et al., 2021).

Collectively, the behavioural, sensor- and source-level findings revealed early disruption of large-scale EM networks in future AD progressors, despite normal cognition at testing, consistent with prior characterisations of subjective cognitive decline (Olichney et al., 2011; Jessen et al., 2020; Musaeus et al., 2019; Paitel et al., 2021; Dubois et al., 2021). Such abnormalities support the progressive disconnection AD syndrome, whereby silent Aβ and tau accumulation disrupts posterior sensory–memory networks and their integration with frontal control regions via synaptic dysfunction and reduced cognitive reserve (Braak and Braak, 1991; Stern, 2012; Palmqvist et al., 2017; Dubois et al., 2018; Busche and Hyman, 2020; Hampel et al., 2021; Dubois et al., 2021). This memory network breakdown, likely increases reliance on compensatory decision strategies, manifesting behaviourally as slower less accurate recognition, mirroring visual-semantic and monitoring deficits and hallmarks of AD’s amnesic hippocampal syndrome (Roper, 2011; Olichney et al., 2011; Friedman, 2013; Musaeus et al., 2019; Paitel et al., 2021).

Together these results highlight the potential of integrated EEG-based cognitive markers for early detection and risk stratification in the preclinical stage of AD, when therapeutic windows remain most effective (Sperling et al., 2011; Hampel et al., 2021; Dubois et al., 2021; Alzheimer’s Association, 2024).

## Contributors Role

The Filipa Raposo Pereira wrote the original draft, edited, curated the data used and was involved in the material preparation, project administration, methodology, formal analysis and investigation, data curation, and visualization. Maximilien Chaumon was involved in the formal analysis, methodology, and writing — review. Bruno Dubois was responsible for the conceptualization, was involved in the funding acquisition, project administration, resources, and writing — review. Hovagim Bakardjian was involved in the conceptualization and investigation and writing — review. Katia Andrade was involved in the writing — review. Marie-Odile Habert was involved in the formal analysis and methodology of PET exams and in the writing — review. Nadjia Younsi was involved in the resources. Valentina La Corte was involved in the conceptualization, methodology, resources, supervision, project management, validation, funding acquisition, and writing — review & editing. Nathalie George was involved in the formal analysis, project managing, methodology, resources, supervision, validation, funding acquisition, and writing — review & editing. All authors read and approved the final manuscript.

## Data Availability

The current study is based on data from the INSIGHT-preAD cohort, whose principal investigator is Prof. Bruno Dubois. Anonymized data are available to all qualified investigators. However, the data from the INSIGHT-preAD study cannot be publicly shared due to privacy and ethical restrictions. Requests to access the data should be sent to Stéphanie Bombois and will be reviewed by the INSIGHT-preAD Scientific Advisory Board. Upon approval, access will be granted via a material transfer agreement with INSERM, considering data sharing restrictions imposed by the local ethics committee and applicable privacy and data protection laws. The code used for data analysis is available at: https://osf.io/kbzdr/overview?view_only=b5b0b7ddbd5d42f3afb1e7e9b03a36c1

## Acknowledgements

Data used in preparation of this article were obtained from the INveStIGation of AlzHeimer’s PredicTors in Subjective Memory Complainers study (INSIGHT-preAD) database. As such, we thank the researchers within the INSIGHT-preAD project who contributed to the experimental design, logistics, implementation of the study, and data collection. We are grateful to Masha Bahrami in particular, for her help in data acquisition. Lastly, we thank all INSIGHT-preAD participants for their commitment to the study.

## Conflicts

The authors have no competing interests. The funding bodies had no role in the study design, in the data analysis and interpretation, or in the article writing.

## Funding Sources

This study was funded by a grant from the Fondation pour la Recherche sur Alzheimer to Nathalie George, Valentina La Corte, and Filipa Raposo Pereira [FRA, project: ePARAD]. The INSIGHT-preAD study [IDRCB: 2012-A01731-42] was legally sponsored by Inserm [protocol nb. C12-59]. It received funding from the program “Investissements d’avenir” [ANR-10-IAIHU-06] and from Pfizer. The Fondation Plan-Alzheimer further funded the MRI and the ¹⁸F-fluorodeoxyglucose PET along with the MEMENTO study. Avid provided the ¹⁸F-florbetapir ligand for Aβ PET. The study was done in collaboration with the Centre Hospitalier Universitaire de Bordeaux [study number CIC EC7].

## INSIGHT-preAD Study Group members

Bonheur J, Boukadida L, Boukerrou N, Cavedo E, Chiesa P A, Colliot O, Dubois B, Dubois M, Epelbaum S, Gagliardi G, Genthon R, Habert M-O, Hampel H, Houot M, Kas A, Lamari F, Levy M, Lista S, Metzinger C, Mochel F, Nyasse F, Poisson C, Potier M-C, Revillon M, Dos Santos A, Santos-Andrade K, Sole M, Surtee M, Thiebaut de Schotten M, Vergallo A, Younsi N, Audrain C, Auffret A, Bakardjian H, Baldacci F, Batrancourt B, Benakki I, Benali H, Bertin H, Bertrand A, Cacciamani F, Causse V, Cherif Touil S, Dalla Barba G, Depaulis D, Fontaine B, Francisque F, Genin A, Glasman P, Gombert F, Hewa H, Jungalee N, Kilani M, La Corte V, Le Roy F, Lehericy S, Letondor C, Lowrey M, Ly J, Makiese O, Masetti I, Mendes A, Michon A, Nait Arab R, Perrin C, Poirier F, Ratovohery S, Rojkova K, Schindler R, Servera M C, Seux L, Simon V, Skovronsky D, Thiebaut M, Uspenskaya O, Vlaincu M, Bombois S, Chupin M, Mangin JF.

## Notes

https://osf.io/kbzdr/overview?view_only=b5b0b7ddbd5d42f3afb1e7e9b03a36c1

### Competing Interest Statement

The authors declare no competing interests directly related to this manuscript. However, the INSIGHT-preAD cohort received partial funding from Pfizer and from Avid Radiopharmaceuticals for cohort-level data acquisition; neither entity had any role in the design, analysis, interpretation, or writing of this specific study.

## References

1. Alzheimer’s Association. 2024 Alzheimer’s disease facts and figures. Alzheimers Dement. 2024;20(5):3708–3821. doi:10.1002/alz.13809

2. Arenaza-Urquijo EM, Vemuri P. Resistance vs resilience to Alzheimer disease: clarifying terminology for preclinical studies. Neurology. 2018;90(15):695–703. doi:10.1212/WNL.0000000000005303

3. Babiloni C, Lopez S, Del Percio C, et al. Resting-state posterior alpha rhythms are abnormal in subjective memory complaint seniors with preclinical Alzheimer’s neuropathology and high education level: the INSIGHT-preAD study. Neurobiol Aging. 2020;90:43–59. doi:10.1016/j.neurobiolaging.2020.01.012

4. Braak H, Braak E. Neuropathological staging of Alzheimer-related changes. Acta Neuropathol. 1991;82(4):239–259. doi:10.1007/BF00308809

5. Busche MA, Hyman BT. Synergy between amyloid-β and tau in Alzheimer’s disease. Nat Neurosci. 2020;23(10):1183–1193. doi:10.1038/s41593-020-0687-6

6. Cabeza R, Albert M, Belleville S, et al. Maintenance, reserve and compensation: the cognitive neuroscience of healthy ageing. Nat Rev Neurosci. 2018;19(11):701–710. doi:10.1038/s41583-018-0068-2

7. Cecchi M, Moore DK, Sadowsky CH, et al. A clinical trial to validate event-related potential markers of Alzheimer’s disease in outpatient settings. Alzheimers Dement (Amst). 2015;1(4):387–394. doi:10.1016/j.dadm.2015.08.004

8. Curran T. Brain potentials of recollection and familiarity. Mem Cognit. 2000;28(6):923–938. doi:10.3758/bf03209340

9. Dubois B, Slachevsky A, Litvan I, Pillon B. The FAB: a Frontal Assessment Battery at bed-side. Neurology. 2000;55(11):1621–1626. doi:10.1212/wnl.55.11.1621

10. Dubois B, Epelbaum S, Nyasse F, et al.; INSIGHT-preAD Study Group. Cognitive and neuroimaging features and brain β-amyloidosis in individuals at risk of Alzheimer’s disease (INSIGHT-preAD): a longitudinal observational study. Lancet Neurol. 2018;17(4):335–346. doi:10.1016/S1474-4422(18)30029-2

11. Dubois B, Villain N, Frisoni GB, et al. Clinical diagnosis of Alzheimer’s disease: recommendations of the International Working Group. Lancet Neurol. 2021;20(6):484–496. doi:10.1016/S1474-4422(21)00066-1

12. Folstein MF, Folstein SE, McHugh PR. “Mini-mental state”. A practical method for grading the cognitive state of patients for the clinician. J Psychiatr Res. 1975;12(3):189–198. doi:10.1016/0022-3956(75)90026-6

13. Friedman D, Johnson R Jr. Event-related potential (ERP) studies of memory encoding and retrieval: a selective review. Microsc Res Tech. 2000;51(1):6–28. doi:10.1002/1097-0029(20001001)51:1<lt;6::AID-JEMT2>3.0.CO;2-R

14. Friedman D. The cognitive aging of episodic memory: a view based on the event-related brain potential. Front Behav Neurosci. 2013;7:111. doi:10.3389/fnbeh.2013.00111

15. Grober E, Buschke H, Crystal H, et al. Screening for dementia by memory testing. Neurology. 1988;38(6):900–903. doi:10.1212/wnl.38.6.900

16. Grober E, Lipton RB, Hall C, Crystal H. Memory impairment on free and cued selective reminding predicts dementia. Neurology. 2000;54(4):827–832. doi:10.1212/wnl.54.4.827

17. Grober E, Veroff AE, Lipton RB. Temporal unfolding of declining episodic memory on the Free and Cued Selective Reminding Test in the predementia phase of Alzheimer’s disease: Implications for clinical trials. Alzheimers Dement 2018;10:161–171. doi:10.1016/j.dadm.2017.12.004

18. Habert MO, Bertin H, Labit M, et al. Evaluation of amyloid status in a cohort of elderly individuals with memory complaints: validation of the method of quantification and determination of positivity thresholds. Ann Nucl Med. 2018;32(2):75–86. doi:10.1007/s12149-017-1221-0

19. Hampel H, Hardy J, Blennow K, et al. The Amyloid-β pathway in Alzheimer’s disease. Mol Psychiatry. 2021;26(10):5481–5503. doi:10.1038/s41380-021-01249-0

20. Horváth A, Szucs A, Csukly G, et al. EEG and ERP biomarkers of Alzheimer’s disease: a critical review. Front Biosci (Landmark Ed). 2018;23(2):183–220. doi:10.2741/4587

21. Hothorn T, Bretz F, Westfall P. Simultaneous inference in general parametric models. Biom J. 2008;50(3):346–363. doi:10.1002/bimj.200810425

22. Jack CR Jr, Knopman DS, Jagust WJ, et al. Hypothetical model of dynamic biomarkers of the Alzheimer’s pathological cascade. Lancet Neurol. 2010;9(1):119–128. doi:10.1016/S1474-4422(09)70299-6

23. Jack CR Jr, Wiste HJ, Weigand SD, et al. Defining imaging biomarker cut points for brain aging and Alzheimer’s disease. Alzheimers Dement. 2017;13(3):205–216. doi:10.1016/j.jalz.2016.08.005

24. Jessen F, Amariglio RE, Buckley RF, et al. The characterisation of subjective cognitive decline. Lancet Neurol. 2020;19(3):271–278. doi:10.1016/S1474-4422(19)30368-0

25. Liang Q, Chen Z, Tang X, Wang XJ. Applications of event-related potentials in Alzheimer’s disease: a systematic review and analysis. Front Aging Neurosci. 2025;17:1513049. doi:10.3389/fnagi.2025.1513049

26. Long S, Benoist C, Weidner W. World Alzheimer Report 2023: Reducing dementia risk— never too early, never too late. Alzheimer’s Disease International. 2023.

27. McNair DM, Kahn RJ. Self-assessment of cognitive deficits. In: Crook T, Ferris S, Bartus R, eds. Assessment in Geriatric Psychopharmacology. New Canaan, CT: Mark Powley; 1983:119–136.

28. Morris JC. The Clinical Dementia Rating (CDR): current version and scoring rules. Neurology. 1993;43(11):2412–2414. doi:10.1212/wnl.43.11.2412-a

29. Musaeus CS, Engedal K, Høgh P, et al. Oscillatory connectivity as a diagnostic marker of dementia due to Alzheimer’s disease. Clin Neurophysiol. 2019;130(10):1889–1899. doi:10.1016/j.clinph.2019.07.016

30. Olichney JM, Taylor JR, Gatherwright J, et al. Patients with MCI and N400 or P600 abnormalities are at very high risk for conversion to dementia. Neurology. 2008;70(19 Pt 2):1763–1770. doi:10.1212/01.wnl.0000281689.28759.ab

31. Olichney JM, Yang JC, Taylor J, et al. Cognitive event-related potentials: biomarkers of synaptic dysfunction across the stages of Alzheimer’s disease. J Alzheimers Dis. 2011;26 Suppl 3:215–228. doi:10.3233/JAD-2011-0047

32. Olichney JM, Pak J, Salmon DP, et al. Abnormal P600 word repetition effect in elderly persons with preclinical Alzheimer’s disease. Cogn Neurosci. 2013;4(3-4):143–151. doi:10.1080/17588928.2013.838945

33. Paitel ER, Samii MR, Nielson KA. A systematic review of cognitive event-related potentials in mild cognitive impairment and Alzheimer’s disease. Behav Brain Res. 2021;396:112904. doi:10.1016/j.bbr.2020.112904

34. Palmqvist S, Schöll M, Strandberg O, et al. Earliest accumulation of β-amyloid occurs within the default-mode network and concurrently affects brain connectivity. Nat Commun. 2017;8(1):1214. doi:10.1038/s41467-017-01150-x

35. Picton TW, Bentin S, Berg P, et al. Guidelines for using human event-related potentials to study cognition: recording standards and publication criteria. Psychophysiology. 2000;37(2):127–152.

36. Roper JC. The neural correlates of retrospective memory monitoring: convergent findings from ERP and fMRI [dissertation]. Provo, UT: Brigham Young University; 2011. https://scholarsarchive.byu.edu/etd/3052

37. Rugg MD, Curran T. Event-related potentials and recognition memory. Trends Cogn Sci. 2007;11(6):251–257. doi:10.1016/j.tics.2007.04.004

38. Razafimahatratra S, Guieysse T, Lejeune FX, et al. Can a failure in the error-monitoring system explain unawareness of memory deficits in Alzheimer’s disease? Cortex. 2023;166:428–440. doi:10.1016/j.cortex.2023.05.014

39. Raposo Pereira F, Chaumon M, Dubois B, et al. Recognition memory decline is associated with the progression to prodromal Alzheimer’s disease in asymptomatic at-risk individuals. J Neurol. 2024b;272(1):70. doi:10.1007/s00415-024-12834-y

40. Raposo Pereira F, George N, Dalla Barba G, et al.; INSIGHT-preAD Study Group. The Memory Binding Test detects early subtle episodic memory decline in preclinical Alzheimer’s disease: a longitudinal study. J Alzheimers Dis. 2024a;98(2):465–479. doi:10.3233/JAD-230921

41. Sperling RA, Aisen PS, Beckett LA, et al. Toward defining the preclinical stages of Alzheimer’s disease: recommendations from the National Institute on Aging–Alzheimer’s Association workgroups on diagnostic guidelines for Alzheimer’s disease. Alzheimers Dement. 2011;7(3):280–292. doi:10.1016/j.jalz.2011.03.003

42. Stern Y. Cognitive reserve in ageing and Alzheimer’s disease. Lancet Neurol. 2012;11(11):1006–1012. doi:10.1016/S1474-4422(12)70191-6

43. Tadel F, Baillet S, Mosher JC, Pantazis D, Leahy RM. Brainstorm: a user-friendly application for MEG/EEG analysis. Comput Intell Neurosci. 2011;2011:879716. doi:10.1155/2011/879716

44. Tsivilis D, Allan K, Roberts J, et al. Old-new ERP effects and remote memories: the late parietal effect is absent as recollection fails whereas the early mid-frontal effect persists as familiarity is retained. Front Hum Neurosci. 2015;9:532. doi:10.3389/fnhum.2015.00532

45. Tzourio-Mazoyer N, Landeau B, Papathanassiou D, et al. Automated anatomical labeling of activations in SPM using a macroscopic anatomical parcellation of the MNI MRI single-subject brain. Neuroimage. 2002;15(1):273–289. doi:10.1006/nimg.2001.0978

46. Van der Linden M, Coyette F, Poitrenaud J, Kalafat M. L’épreuve de rappel libre / rappel indice à 16 items (RL/RI-16). In: Van der Linden M, Adam S, Agniel A, Baisset Mouly C, et al., editors. L’évaluation des troubles de la mémoire: présentation de quatre tests de mémoire épisodique (avec leur étalonnage). Solal Editeur; 2004.

47. Wang Y, Li Q, Yao L, et al. Shared and differing functional connectivity abnormalities of the default mode network in mild cognitive impairment and Alzheimer’s disease. Cereb Cortex. 2024;34(3):bhae094. doi:10.1093/cercor/bhae094.

48. Zamani J, Vahid A, Avelar-Pereira B, et al.; Alzheimer’s Disease Neuroimaging Initiative. Mapping amyloid beta predictors of entorhinal tau in preclinical Alzheimer’s disease. Alzheimers Dement. 2025;21(2):e14499. doi:10.1002/alz.14499

