## supplementary material for "Episodic memory ERPs reflect preclinical Alzheimer’s disease progression"

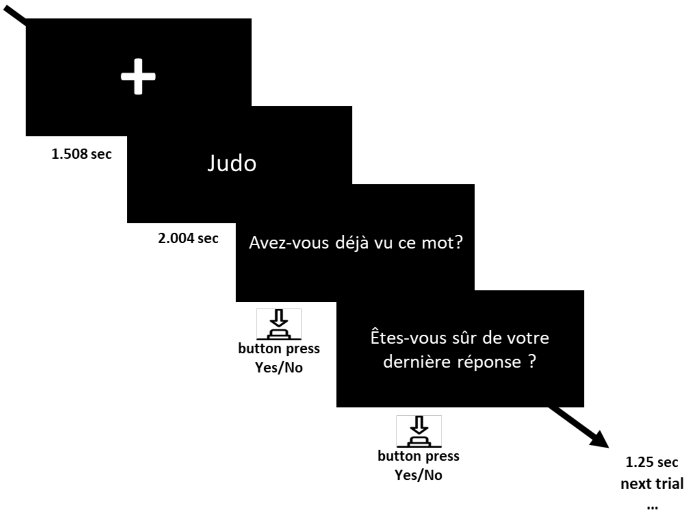


**Supplementary Figure 1. Sample trial of the Old/New word recognition task.** Translation: “Avez-vous déjà vu ce mot?” means “Did you see this word before?”; “Etes-vous sûr de votre dernière response?” means “Are you sure of your response?”.

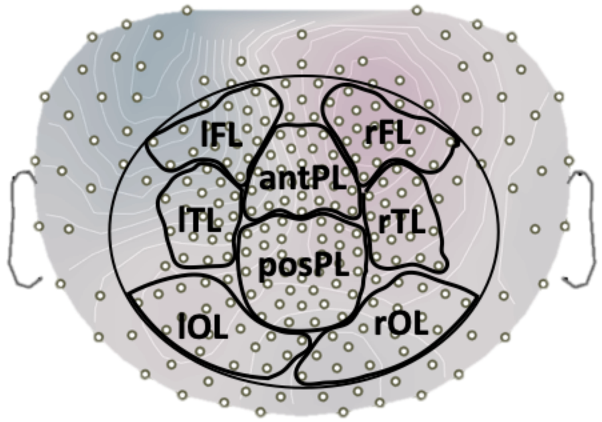


**Supplementary Figure 2. Regions of interest (ROIs) defined on the EEG sensor layout.**
The figure shows a standard EEG GSN HydroCel 256 montage with ROIs defined over scalp areas most consistently implicated in Old/New event-related potentials (ERP) effects during episodic memory retrieval. The black circle delineates the subset of electrodes included in the analysis. Left and right hemispheres, and anterior and posterior regions were defined symmetrically, each containing the same number of electrodes. The selected ROIs were: left frontal lobe (lFL), right FL (rFL), left temporal lobe (lTL), right TL (rTL), anterior parietal lobe (antPL), posterior PL (posPL), left occipital lobe (lOL), and right OL (rOL).

**Supplementary Table 1. Number of electrodes included on each region of interest (ROI).** The numbers in the table indicate the electrode numbering labels according to the EGI GSN hydrocel 256 system. The number of electrodes was defined symmetrically per hemisphere and between the anterior posterior regions as follows:  left frontal lobe (lFL), right FL (rFL), left temporal lobe (lTL), right TL (rTL), anterior parietal lobe (antPL), posterior PL (posPL), left occipital lobe (lOL), and right OL (rOL).

| ROIs | Electrode numbers on each ROI |
|---|---|
| **lelft FL** | 28; 29; 30; 34; 35; 36; 39; 40; 41; 42; 48; 49; 50; 51; 55, 56; 61; 62 |
| **right FL** | 2; 3; 45; 13; 196; 205; 206; 211; 212; 213; 214; 215; 220; 221; 222; 223; 224 |
| **left TL** | 57; 58; 59; 63; 64; 65; 66; 69; 70; 71; 72; 74; 75; 76; 77; 84; 85; 86 |
| **right TL** | 163; 164; 171; 172; 173; 179; 180; 181; 182; 183; 191; 192; 193; 194; 195; 202; 203; 204 |
| **left OL** | 147; 148; 149; 157; 158; 159; 160; 161; 167; 168; 169; 170; 176; 177; 178; 189; 190 |
| **right OL** | 94; 95; 96; 97; 104; 105; 106; 107; 113; 114; 115; 122; 123; 124; 135; 136; 137 |
| **anterior PL** | 6; 7; 8; 9; 15; 16; 17; 23; 24; 43; 44; 45; 52; 53; 60; 81; 132; 144; 155; 184; 185; 186; 197; 198; 207; Cz |
| **posterior PL** | 78; 79; 80; 87; 88; 89; 90; 98; 99; 100; 101; 108; 109; 110; 116; 117; 118; 119; 125; 126; 127; 128; 129; 130; 131; 138; 139; 140; 141; 142; 143; 150; 151; 152; 153; 154 |

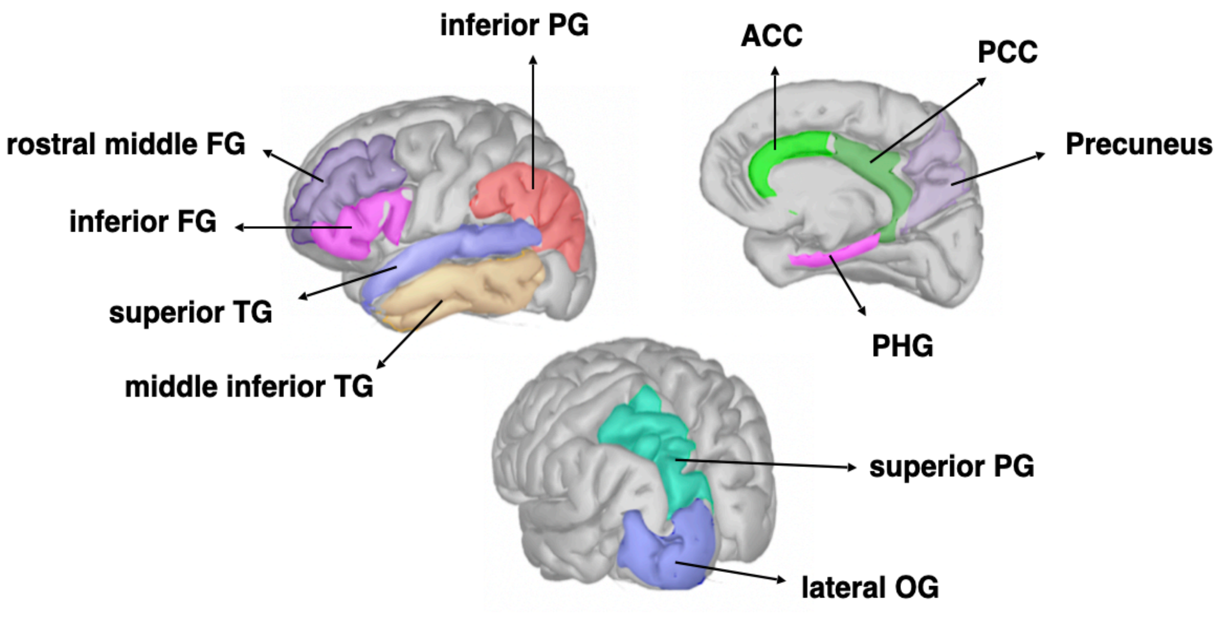


**Supplementary Figure 3. Anatomical regions of interest (ROIs) selected from the Automated Anatomical Labelling (AAL) atlas.**

The figure shows the sagittal, medial sagittal and posterior views of the brain coloured differently according with the ROIs chosen for source analysis based on their known relevance to episodic memory processes and Alzheimer's disease. Several ROIs represent merged AAL regions with nomenclature adapted to better reflect functionally coherent structures. The selected ROIs (considerer bilaterally) included: rostral middle Frontal Gyrus (FG); inferior FG (pars orbitalis, triangularis, and opercularis); superior Temporal Gyrus (TG); middle-inferior TG (middle TG, superior temporal sulcus, and inferior TG); lateral Occipital Gyrus(OG); superior Parietal Gyrus(PG);  Parahippocampal Gyrus (PHG; parahippocampus and entorhinal); precuneus; posterior Cingulate Cortex (PCC; posterior cingulate and isthmus); anterior CC (ACC; anterior rostral cingulate and anterior caudal cingulate); inferior PG (angular gyrus and temporal-parietal junction).

**Supplementary Table 2. Number of subjects considered on each session per group (controls, stables, progressors)**

| | Groups | | |
|---|---|---|---|
| | **Controls** | **Stable** | **Progressors** |
| **M0** | 15 | 15 | 15 |
| **M12** | 15 | 14 | 15 |
| **M24** | 15 | 14 | 14 |
| **M36** | 13 | 12 | 11 |
| **M48** | 15 | 12 | 9 |
| **M60** | 14 | 12 | 8 |

**Supplementary Table 3. ROIs showing significant differences in mean amplitude (±SD; in µV) in the time-windows corresponding to the ERPs of interest—P3 (252–340 ms), FN400 (412–572 ms), P600 (620–772 ms), and post-retrieval monitoring (872–1040 ms)—during the performance of the Old/New word recognition task.**

| ERP | Hemisphere | Selected clusters | Word-Category | | | |
|---|---|---|---|---|---|---|
| | | | OLD | NEW | | |
| | | | mean ± SD | mean ± SD | F(df1, df2) = value | *p* |
| P3 | LH | Frontal | 1.84 ± 0.4 | 0.99 ± 0.4 | F(1, 6.2) = 37.0 | 0.0008 |
| | RH | Frontal | -24.0 ± 0.5 | -1.70 ± 0.4 | F(1, 8.3) = 28.8 | 0.0006 |
| FN400 | LH | Frontal | 1.02 ± 0.5 | -0.29 ± 0.5 | F(1, 7.9) = 34.9 | 0.0004** |
| | | Temporal | 1.34 ± 0.4 | 0.58 ± 0.3 | F(1, 6.7) = 19.7 | 0.0003** |
| | RH | Frontal | -0.30 ± 0.5 | 0.62 ± 0.5 | F(1, 9.4)= 25.5 | 0.0006** |
| | | Temporal | -1.35 ± 0.3 | -0.87 ± 0.3 | F(1, 7.9) = 6.92 | 0.03* |
| | | Occipital | -0.24 ± 0.4 | -0.19 ± 0.4 | F(1, 10.4) = 7.47 | 0.02* |
| | | anterior Parietal | 1.81 ± 0.3 | 1.34 ± 0.3 | F(1, 3.7) = 43.7 | 0.004** |
| P600 | LH | Frontal | -1.93 ± 0.5 | -2.77 ± 0.4 | F(1, 10.8) = 7.0 | 0.02* |
| | | Temporal | 0.62 ± 0.4 | -0.98 ± 0.3 | F(1, 9.2) = 39.0 | 0.0001*** |
| | | Occipital | -0.07 ± 0.5 | -0.12 ± 0.4 | F(1, 14.9) = 10.1 | 0.006** |
| | RH | Frontal | 0.21 ± 0.6 | 2.58 ± 0.4 | F(1, 11.5)= 69.7 | <0.0001*** |
| | | posterior Parietal | 4.03 ± 0.6 | 1.88 ± 0.5 | F(1, 10.1) = 47.8 | <0.0001*** |
| Post-retrieval monitoring | LH | Frontal | -2.09 ± 0.5 | -0.33 ± 0.4 | F(1, 13.6)= 17.1 | 0.001** |
| | | Temporal | -0.19 ± 0.3 | -1.34 ± 0.3 | F(1, 10.5) = 24.1 | 0.0005** |
| | RH | Frontal | 1.11 ± 0.6 | 0.33 ± 0.6 | F(1, 19.0) = 70.4 | <0.0001*** |
| | | posterior Parietal | 2.31 ± 0.6 | 1.17 ± .05 | F(1, 17.8) = 10.2 | 0.005** |

Abbreviations: SD = Standard Deviation; df = degrees of freedom; LH = left hemisphere; RH =right hemisphere. The table represents the mean amplitude (±SD) for Old and New words in clusters and time-windows corresponding to the ERPs of interest (P3, FN400, P600, post-retrieval monitoring) where significant differences were found longitudinally between word-category (Old/ New) as a product of linear mixed effect models (lmer). Significance is based on the following p-values: ***, $p < 0.001$; **, $p \leq 0.01$; *, $p \leq 0.05$.
